# Interspecies exchange of a mobile genetic element during a plant disease outbreak

**DOI:** 10.1101/2024.10.16.618685

**Authors:** Daria Evseeva, Yann Pecrix, Marek Kucka, Catrin Weiler, Carolin Franzl, Markéta Vlková-Žlebková, Elena Colombi, Yingguang Frank Chan, Stéphane Poussier, Emmanuel Wicker, Honour McCann

## Abstract

The evolutionary processes underlying disease outbreaks remain unknown for most bacterial plant pathogens. We sequenced an outbreak of lethal wilt disease in Martinique caused by two distantly related lineages in the *Ralstonia solanacearum* species complex. One lineage (*R. solanacearum* IIB-4NPB) exhibited the broadest host range ever documented for a single lineage in the field, while the other (*R. pseudosolanacearum* I-18) expanded in parallel but retained specialisation on solanaceaous hosts. Phylogenomic analysis of 407 outbreak isolates shows both lineages independently disseminated from mainland populations into Martinique. We resolved fine-scale geographic patterns of genomic diversity and identified spatial hotspots of interspecies mobile element exchange, resulting in the identification of a new family of *Ralstonia* integrative conjugative elements (ICEs) associated with the outbreak. ICE accessory gene integration sites display striking functional specialization and differentiation despite variable gene content: each site acquires only metabolism-associated or defence-related genes, respectively. This work provides insight into the origin and genomic changes associated with an outbreak of plant disease, and highlights the role of mobile elements in driving pathogen emergence.

---

Disease outbreaks on crop plants have severe consequences for subsistence and commercial growers alike, yet we have little understanding of the genetic mechanisms and evolutionary processes underlying pathogen emergence and adaptation to agricultural environments. Whole genome sequencing of emerging and established bacterial pathogens has led to significant new discoveries detailing the origins of outbreak lineages, dynamic selection pressure imposed by viral predation, and the role of mobile elements in the movement of antimicrobial resistance and virulence genes^1–4^. Despite recent notable exceptions, most studies focus on mammalian pathogens and it is unclear how generalizable their findings are to plant pathogens^5–10^. Diseases can spread quickly across vast areas used for food, forage and industrial crops, with severe consequences for agricultural productivity and economic stability. Clarifying the mechanisms underlying disease emergence in plant hosts is of both fundamental importance and practical value, and deepens our understanding of how microbes adapt to new hosts and environments.

The identification of a novel lineage of *Ralstonia solanacearum* in the Caribbean island of Martinique represents a rare opportunity to investigate the emergence of a bacterial plant pathogen of global economic importance^11^. Members of the *R. solanacearum* species complex (RSSC) are known to cause lethal infections in many different host species, including staple and commodity crops^12–14^. The species complex is comprised of *R. solanacearum* (Rso, also known as phylotype II and further subdivided into Rso IIA and IIB), *R. pseudosolanacearum* (Rps, phylotype I and III), and R. syzygii (Rsy, phylotype IV)^15,16^. Lineages identified within each species or phylotype were assigned to over 56 sequevars using sequence typing of the endoglucanase (*egl*) gene^17,18^. Since the pathogen disseminates rapidly and can persist for years in freshwater and soil, the pathogen is subject to extensive monitoring and movement controls to limit its spread. Clonally propagated staples like potato and banana are particularly threatened by emergence events, since pathogens can spread rapidly in areas with high host density and genetic homogeneity^19,20^. The threat posed to potato cultivation is so great that one Rso lineage is classified as a Select Agent in the United States^21^. Multiple lineages of *R. solanacearum* are known to infect banana, which is cultivated on an industrial scale in the Americas^8^. An outbreak of disease on a cultivated ornamental (*Anthurium andreanum*) in Martinique in 1999 caused alarm when it was found to be caused by a lineage closely related to the banana-infecting lineage Rso IIB sequevar 4 (IIB-4)^11^. Additional infections on the island were soon reported in many hosts which *R. solanacearum* had not been previously known to infect. The threat posed to banana cultivation and apparent host range expansion led to an extensive sampling survey of *Ralstonia* populations across Martinique, as well as coastal and inland areas of French Guiana on the South American mainland^23,24^. This work generated a remarkable collection of strains isolated from infected crops, asymptomatic weeds, soil and freshwater environments during a disease emergence event characterized by the rapid and unprecedented expansion of *Ralstonia* spp. infections across a broad range of hosts and geographic areas in Martinique^11^. Multi-locus sequence analysis (MLSA) showed both Rps and Rso isolates were recovered, but the origins, diversity and genomic changes accompanying the emergence of the host-range expanded lineage remain unclear. Since this collection provides a unique opportunity to explore the disease origins and investigate the spatial population structure of isolates collected from cultivated crops, wild hosts and environmental sources, we sequenced the largest collection of *Ralstonia* spp. ever sampled during an outbreak event.

Our study of 409 new genomes, paired with 407 publicly available genomes, provides unparalleled phylogenetic and spatial resolution of an extraordinarily well sampled outbreak of *Ralstonia* spp. in the Caribbean island of Martinique. Earlier MLSA studies showed two different lineages (Rso IIB-4NPB and Rps I-18) were circulating during the outbreak event; we show here each lineage was disseminated from well-established populations on the South American mainland, though one lineage is of more distant Asian origin and retains a signature of host specialisation. The lineage that underwent an apparent host range expansion arose from a population that has been diversifying in the Americas for much longer. Although each lineage is a member of a different species, we found mobile elements called integrative and conjugative elements (ICEs) are circulating between them, particularly in geographic areas where lineages co-occur. The ICEs have two cargo gene insertion hotspots that exhibit consistent functional differentiation: one hotspot is a target for metabolic gene acquisition, while the other is a hotspot for defense element acquisition. In contrast, there were very few changes in the repertoire of secreted virulence proteins linked with pathogenesis and host defense manipulation. These results challenge the view plant pathogen emergence is driven by virulence factor acquisition or innovation, highlighting instead the critical roles of horizontal transfer, microbial competition and metabolic flexibility in shaping apparent host range and ecological success. The expansion of Rso IIB-4NPB in particular, which exhibits the broadest documented host range of any Ralstonia lineage, was more strongly driven by the acquisition of mobile elements and expanded anti-phage defences than by virulence gene acquisition. As agricultural intensification continues to reshape plant-microbe interactions, pathogen lineages that can rapidly acquire and integrate novel genetic material may rapidly evolve to exploit novel niches and hosts. Surveillance strategies should therefore consider not just the pathogen, but the community and network of genetic exchange that can fuel its evolution.

## Results

### Global phylogeography of the *Ralstonia solanacearum* species complex

The global phylogeography of the *Ralstonia solanacearum* species complex (RSSC) reveals patterns of long-distance dissemination and epidemic expansion. Our analysis of 409 new whole genome sequences from *Ralstonia* isolated from crops, weeds, soil and freshwater environments during the outbreak paired with 407 genomes available on NCBI shows that diverse populations of both *R. solanacearum* (Rso) and *R. pseudosolanacearum* (Rps) are circulating in the Americas ^11,23,24^. The outbreak documented in Martinique was caused by the expansion of two lineages from each species: Rps I-18 and Rso IIB-4NPB (Fig. 1, Fig. S1), consistent with the MLSA results in Wicker et al. 2007. The center of origin of Rso and Rps is in the Americas and Asia, respectively^12^. Multiple inter-continental dissemination events of both Rso and Rps lineages are evident in the tree, including a global outbreak of Rso IIB-1 on potato (*S. tuberosum*) and multiple intercontinental introductions of at least eight distinct Rps lineages from Asia into the Americas (Fig. 1, Fig. S1). Figure 1 shows the Rps lineages introduced to the Americas are divergent (with the exception of Rps I-14 and I-18) and provides phylogenetic context for their distinct origins, placing them in the context of the diversity of Asian Rps. The introduction of Rps lineages into the Americas resulted in their establishment and local diversification, primarily in solanaceous hosts.

**Fig. 1.**
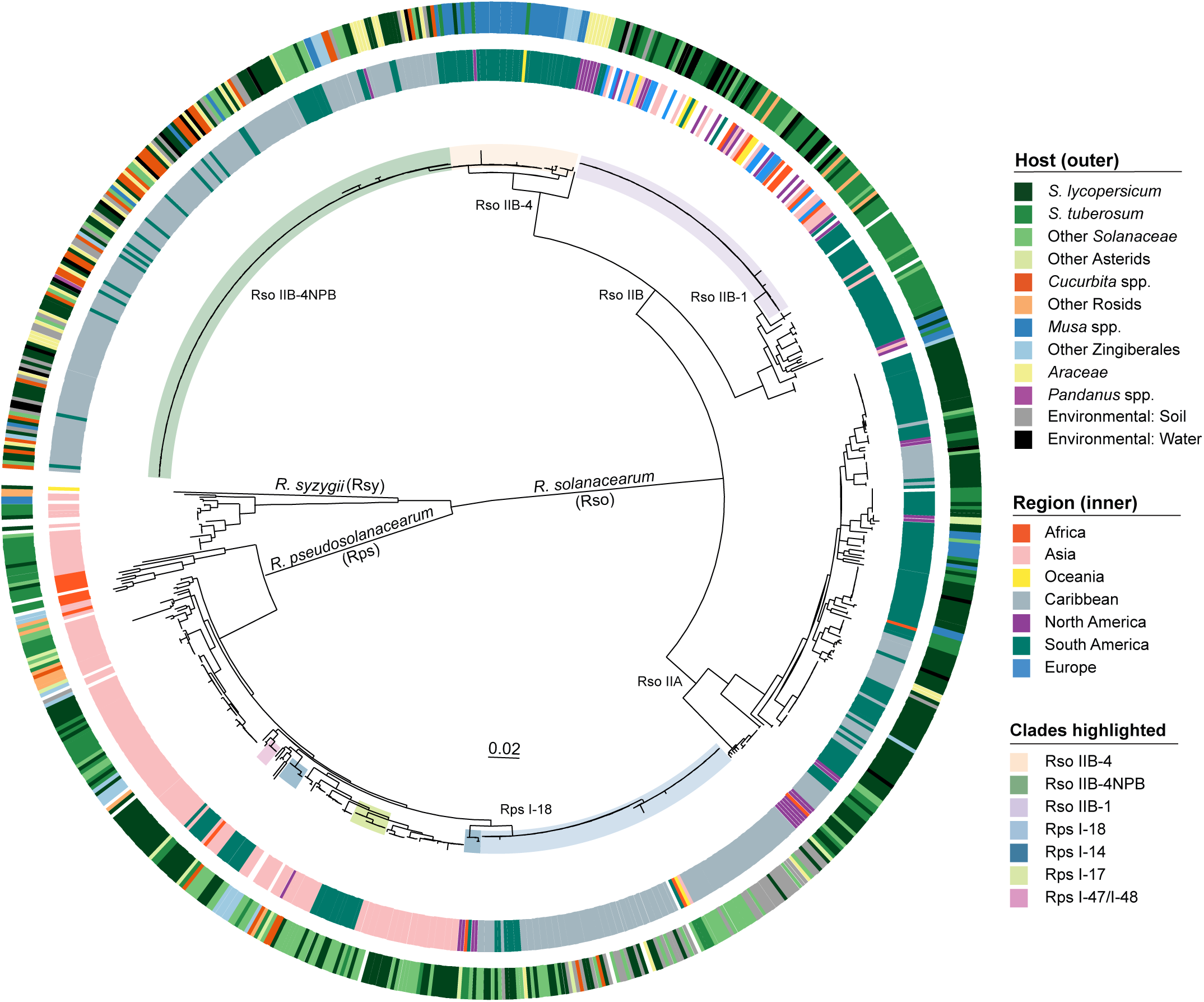
Core gene tree of the *Ralstonia solanacearum* species complex. Maximum likelihood tree of 816 RSSC genomes inferred from a 137,423bp SNP alignment obtained after core gene identification, alignment generation and invariant site removal. Nodes with the bootstrap support values under 70 are collapsed. Clades representing Rso and Rps lineages of interest are highlighted in the tree. The geographic region and host of isolation are shown in the first and second rings, respectively (A and B). Martinique isolates are a subset of Caribbean isolates, and French Guiana isolates are a subset of South American isolates. With the exception of *S. lycopersicum* (tomato) and *S. tuberosum* (potato), the host of isolation for each isolate is shown collapsed to family level. Lower level taxonomic classifications within the same family are indicated with greater colour saturation. Missing data is shown in white.

Rps I-18 is widely distributed across Martinique, French Guiana, Brazil and Peru (Fig. 1, Fig. S1). This lineage shares a recent common ancestor with Rps I-14, which is present in Guatemala, Costa Rica and Colombia. At least three additional Rps lineages of Asian origin are present in the Americas: Rps I-13, Rps I-14, Rps I-17 and Rps I-47/48. Although a single dissemination event could be responsible for the introduction of a diverse pathogen population, it is more likely multiple independent introductions of Rps occurred. The presence of Rps I in the Americas has been documented as early as 1967^25^, yet the genomic epidemiology work shown here provides the first clear evidence of the distinctly different origins and fates of multiple intercontinental dissemination events from Asia into the Americas.

In contrast to Rps, Rso has been diversifying in the Americas for some time. Although two lineages (Rso IIB-4NPB and Rps I-18) expanded in Martinique and French Guiana, it is the Rso IIA isolates that encompass the greatest phylogenetic diversity. Rso IIA, primarily isolated from tomato (68%), is broadly distributed across the Americas, though there is no evidence of recent epidemic expansion (Fig. 1, Fig. S1). Rso IIB lineages are responsible for both regional and global outbreak events however, most notably on banana and potato. Rso IIB-1 (formerly race 3 biovar 2 or R3bv2), responsible for a global outbreak on potato, has also been isolated from other *Solanum* spp., *Pelargonium* spp. (formerly *Geranium* spp.), surface water and soil. Although the centre of origin of Rso IIB-1 appears to be in South America, its rapid global expansion suggests international trade in seed potatoes is the likeliest mechanism of long-distance dissemination, highlighting the elevated risk of disease transmission during movement of infected plant material^14,19,26,27^. Outbreaks of Rso on banana are well documented in the Americas, where it is referred to as Moko disease. Moko disease is clearly polyphyletic, as outbreaks were caused by four Rso IIA and three Rso IIB lineages, including basal members of Rso IIB-4 (Fig. 1)^28,29^. These banana-infecting lineages are responsible for multiple regional outbreaks in Panama, Costa Rica, Colombia, Peru, Brazil and French Guiana from at least the 1960s onwards (Fig. S2). Since banana was only introduced to the Americas as a crop in the 16th century^30^, it is a matter of some interest that the most deeply branching basal members of Rso IIB are isolated from *Musa* spp. along with *Heliconia* spp. and *Anthurium* spp., both endemic to the Americas (Fig. 1, Fig. S2)^31,32^. The expansion of high-density single variety banana monocultures likely increased opportunities for the pathogen to emerge and spread in agricultural areas, in turn increasing likelihood of spread to other crop varieties, both endemic and introduced.

### Epidemic expansion of two lineages after their dissemination from South America

The species complex tree confirms the surge in plant infections in Martinique is due to the recent clonal expansion of two lineages across the island: Rso IIB-4NPB and Rps I-18 (Fig. 1). Rso IIB-4NPB is of particular interest since it was isolated from the most diverse group of naturally infected hosts that has ever been documented for *Ralstonia* to date. Rps I-18 expanded in tandem with Rso IIB-4NPB, however members of this lineage originally only caused infections of solanaceous hosts in the field^11,23^. We investigated the origins of both lineages using well resolved, non-recombinant core genome trees including closely related strains from mainland populations (Fig. 2).

**Fig. 2.**
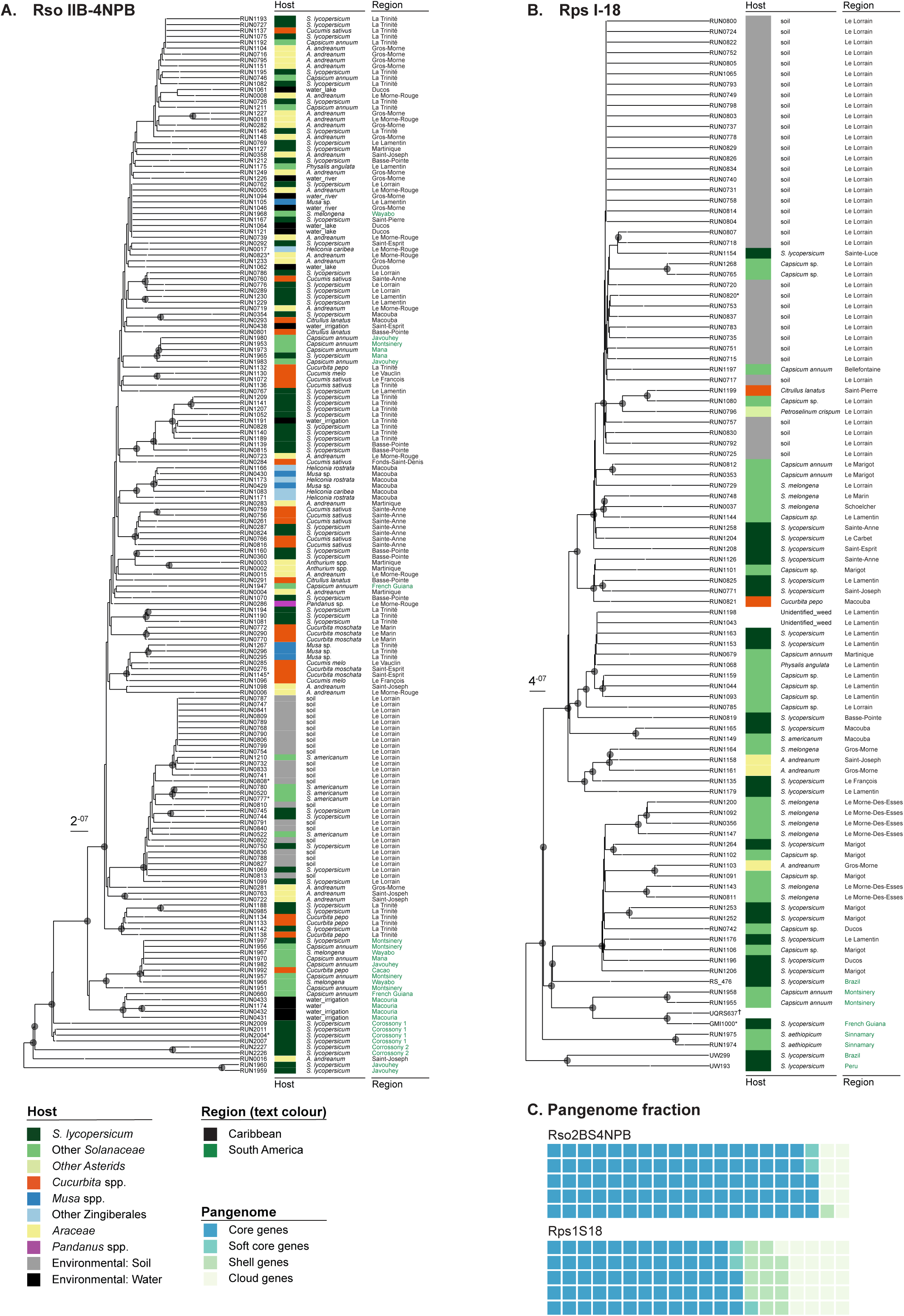
Outbreak lineage core genome trees. (A) Rso IIB-4NPB maximum likelihood tree calculated from a 5,206,677 bp non-recombinant non-mobile core genome alignment with 195 SNPs and 73 parsimony informative sites. Internal nodes with bootstrap value above 80 are indicated with black circles. Strain labels shown in bold are are complete or near complete assemblies generated in this study and/or used as mapping references. (B) Rps I-18 maximum likelihood tree calculated from a 4,760,272 bp non-recombinant core genome alignment with 311 SNPs and 116 parsimony informative sites. The dagger for UQRS637 indicates this assembly was later suppressed on NCBI by the uploader. Asterisks next to strain labels indicates closed genomes generated in this study and/or used as mapping references. (C) Pangenome waffle chart representing the proportion of core (present in 99 - 100% of genomes), soft-core (95 - 99% of genomes), shell (15 - 95% of genomes) and cloud genes (present in less than 15% of genomes) identified by Panaroo, displayed separately for each lineage pangenome.

Basal members of the Rso IIB-4NPB tree were mostly isolated from solanaceous hosts or freshwater sources on the South American mainland, in French Guiana (Fig. 2, Fig. S3). The basal group of isolates likely represents the ancestral mainland population from which Rso IIB-4NPB emerged to sweep across Martinique, since only one of these was isolated on Martinique (RUN0016). There was limited spread of the derived clade on the mainland: only six were isolated from solanaceous plants in French Guiana while the remaining 149 were isolated in Martinique during the epidemic. Although a mainland origin for Rso IIB-4NPB (and indeed Rps I-18) is most likely, it is possible there are more closely related ancestral populations that remain unsampled. The derived group of 155 isolates exhibits limited genetic diversity, differing by only 166 nonrecombinant SNPs in the core genome. Curiously, both basal and derived members of Rso IIB-4NPB were isolated from *Anthurium andreanum* in Saint Joseph, Martinique, suggesting this location and host may be linked with the initial introduction of the emergent lineage to the island (Fig. 2). Notably, Rso IIB-4NPB was also isolated from symptomatic banana in Trinité, indicating the appellation ‘Non-Pathogenic on Banana’ (NPB) for the lineage may not be wholly accurate.

The evolutionary history of Rps I-18 parallels that of Rso IIB-4NPB: Martinique Rps I-18 is similarly derived from a broadly distributed population on the mainland (Fig. 2, Fig. S4). Despite the large number of sequenced Rps I-18 isolates, the lineage displays limited diversity in the core genome and T3E repertoire, consistent with recent epidemic expansion. In contrast to Rso IIB-4NPB, there is no evidence of expanded host range: with few exceptions, the majority of Martinique Rps I-18 was isolated from soil or solanaceous hosts (Fig. S5). There is evidence of spatial population structure for Rps I-18 and Rso IIB-4NPB alike: Le Lorrain Rso IIB-4NPB and Rps I-18 isolates form distinct and well supported clades in their respective trees, regardless of whether they were isolated from soil, cultivated or wild *Solanum* spp.

We used these well-resolved non-recombinant trees to determine the divergence time of Rps I-18 and Rso IIB-4NPB. BactDating showed there is some correlation between the sampling date and root-to-tip distance for Rso IIB-4NPB (R^2^=0.16, p<0.0001) and permutation analysis showed the estimated divergence time outperformed random sampling in 77.6% of simulations. Though weakly supported, the estimated divergence time for the split between the basal mainland and derived Martinique population of Rso IIB-4NPB is between 1930 -1970 (Fig. S6). Rps I-18 exhibits little temporal signal (R^2^=0.05, p=0.04), yet date permutation analysis showed the estimated divergence time outperformed random sampling in 100% of simulations. The estimated divergence time for Rps I-18 is earlier, between 1870 - 1940 (Fig S7). During pangenome analyses we found both Rps I-18 and Rso IIB-4NPB have similar numbers of core and soft-core genes (4,565 for Rps I-18 and 4,792 for Rso IIB-4NPB), but different accessory genome sizes: 487 accessory genes in Rso IIB-4NPB and 2,323 in Rps I-18. While it is possible the difference in accessory genome sizes (amounting to 10% and 35% of the Rso IIB-4NPB and Rps I-18 pangenome, respectively) may be caused by differences in the rate or maintenance of horizontal acquisition events, this may also reflect different introduction times to Martinique. The data cumulatively indicates Rps I-18 was introduced to Martinique prior to Rso IIB-4NPB.

### Ancestral changes in virulence and metabolic gene repertoires

Despite the reduced genetic diversity and recent emergence of Rso IIB-4NPB, derived members of this lineage have been isolated from the widest variety of hosts during natural infections and display the broadest documented host range of any single lineage in the species complex (Fig. 2). Since the delivery of virulence proteins by the Type 3 secretion system is known to play an important role in successful host colonisation and the repertoire of Type 3 secreted effectors (T3Es) in turn is thought to play an important role in host specificity^33–35^, we identified the T3E repertoire across the entire dataset to determine whether the gain or loss of specific T3Es is linked with the emergence of Rso IIB-4NPB (Fig. 3, Fig. S1, Table S4). Rso IIB-4NPB genomes display nearly uniform T3E repertoires, consistent with the recent expansion of this lineage, with no lineage-specific T3E changes compared to Rso IIB-4 (Fig. S2). Multiple T3E changes occurred in both the common ancestor of Rso IIB-4NPB and Rso IIB-4, and in Rso IIB-4 alone. For example, a single T3E is uniquely present in Rso IIB-4 and Rso IIB-4NPB, but absent in all other RSSC genomes: the hypothetical effector *T3E_Hyp13* (UW163_19700 in Rso IIB-4 UW163, OFBKGH_20220 in Rso IIB-4NPB RUN1145) (Fig. S1, Table S6). Another T3E (*ripAX1*) is sufficiently divergent from other members of the effector family to be classified as a clade-specific gene by Panaroo pangenome analysis. Two effectors are present in Rso IIB-4 and absent in Rso IIB-4NPB: *ripAX2* and *ripP3 (*(Fig. S1, Fig. S2, Table S6). The effector *ripAX2* has a broad distribution across Rso IIA and IIB genomes and was therefore likely lost in Rso IIB-4NPB, while *ripP3* has a patchy distribution and was likely acquired by Rso IIB-4. Four effectors appear to have been lost in basal Rso IIB-4: *ripAQ*, *ripAW*, *ripBD* and the predicted *T3E_Hyp3* effector. The loss of *ripAQ* is most striking as the family is otherwise present in all other RSSC strains. Since these effectors are present among related clades and/or across the species complex, the most parsimonious explanation for the pattern observed is they were lost in the ancestor of banana-infecting Rso IIB-4 strains rather than being acquired by ancestral Rso IIB-4NPB. In sum, the Rso IIB-4 isolated from monocot hosts (*Heliconia* spp., banana and *Epipremnum* spp.) exhibit more variation in effector repertoire than Rso IIB-4NPB (Fig. 3, Fig. S2) and contain more unique variants statistically associated with the host of isolation (Supplementary Results p.16).

**Fig. 3.**
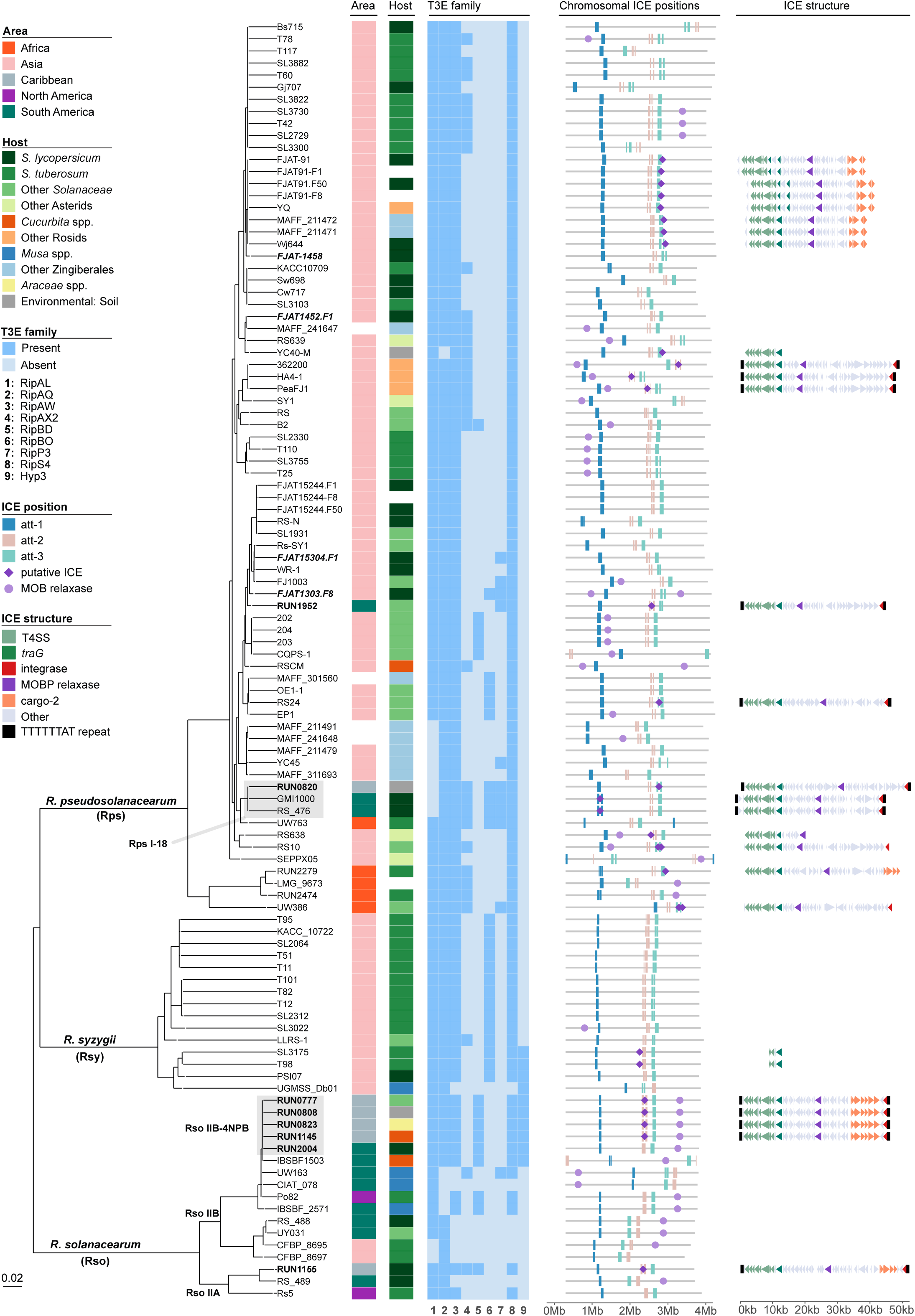
Virulence gene and ICE distribution across the *R. solanacearum* species complex. Maximum likelihood tree of all completely assembled RSSC genomes available in this study (n=105). Strain labels shown in bold are complete or near complete assemblies generated in this study and/or used as mapping references. Strain labels shown in boldface italics represent clades collapsed due to low diversity. The geographical area and host of isolation are shown next to strain labels, followed by nine *Ralstonia* T3E families exhibiting presence/absence variation for Rso IIB-4 and Rps I-18 clades. Dark blue in the T3E family column represents the presence of the T3E family in a strain and light blue represents its absence. Chromosomal ICE position displays the position of the ICE attachment sites on the main chromosome of each RSSC genome – no ICEs were identified on the megaplasmid in any complete genome. The attachment sites are here defined as regions syntenic to the 50kb up- and downstream flanking regions of the Rps I-18 GMI1000 (att-1, blue), Rso IIB-4NPB RUN1145 (att-2, pink) and Rso IIB-4NPB RUN1960 (att-3, green) attachment sites. Thin grey lines represent the length ranges of the chromosomes. MOB relaxases occurring in isolation (purple circles) are separately indicated from those adjacent to T4SS genes and forming putative ICEs (purple diamonds). The final panel shows the organization of putative ICEs identified in RSSC strains, with homologs of various functional categories highlighted in different colours. RS10 and UW386 both have two ICEs inserted in att-3. Black bars around ICEs represent the TTTTTTAT repeat motif delimiting putative ICE boundaries.

We investigated non-T3E acquisitions in the ancestor of Rso IIB-4NPB, and found 84 genes uniquely acquired in the ancestor of both Rso IIB-4 and Rso IIB-4NPB (Table S3). These genes encode enzymes such as a TauD/TfdA family dioxygenase, shown to degrade phenoxy herbicides in soil-borne bacteria like *Burkholderia* spp. and *Cupriavidus necator* (formerly *Ralstonia eutropha*)^36–41^; a CxxC motif protein (UW163_01710, OFBKGH_14170) associated with thiol-disulfide oxidoreductase activity and oxidative damage responses^42,43^; an electron acceptor flavodoxin (UW163_18870, OFBKGH_23185) also involved in oxidative damage response^44^; and a carboxymuconolactone decarboxylase (UW163_18885, OFBKGH_23170) in the protocatechuate catabolism pathway, linked with the degradation of both lignin precursors^45^ and the conversion of lignin-derived aromatic compounds to tricarboxylic acid intermediates like succinate and malate^46,47^. In sum, although the function of many horizontally acquired genes remains unclear, some ancestral acquisitions appear to be linked with the catabolism of aromatic compounds in soil or plant tissues.

### Integrative and conjugative elements circulating between co-occurring pathogen lineages

Since the co-occurrence of two pathogen lineages in the same host or niche provides opportunities for homologous recombination, we then explored whether Rps I-18 and Rso IIB-4NPB were exchanging genetic material. We found no evidence of homologous recombination between Rso IIB-4NPB and Rps I-18 (Fig. S8). RECOPHY identified some regions with reduced (< 10/kb) and elevated (120/kb) SNP density compared to the alignment-wide average of 53 SNPs/kb, yet distance trees for those regions display the same topology as the core genome tree, indicating homologous recombination between these lineages is unlikely to have occurred^47,48^. ClonalFrameML analysis indicated that although the ratio of recombination to mutation in both lineages is low (R/theta = 0.159 for Rps I-18 and 0.152 for Rso IIB-4NPB), homologous recombination is occurring with unknown donors.

Mobile elements constitute another means by which genetic material can be exchanged between bacterial pathogens. Pangenome exploration and association testing alike revealed the presence of integrative and conjugative elements (ICEs) present in co-occurring Rps I-18 and Rso IIB-4NPB, but absent in Rso IIB-4 (Fig. 3). All but one of the 27 genes associated with isolation in Martinique by SCOARY are located on an ICE, while a PySEER kmer-based association test similarly showed 625 out of 2,367 Martinique-associated kmers and all 83 soil-associated kmers map to the same ICE (Fig. S9, Table S10, Table S11, Table S12). A clear association between isolation from Trinité and Morne Rouge and a cluster of eight ICE genes was identified, and another two ICE gene clusters display overlapping associations with isolation from soil and Le Lorrain (Fig. S9, Fig. S10, Fig. S11). The gene clusters linked with specific locations of isolation in Martinique correspond to ICE backbone genes and cargo gene modules .

Since the emergent Rso IIB-4NPB lineage acquired an ICE absent in Rso IIB-4, we predicted ICEs in all 133 complete *Ralstonia* genomes (Fig. 3). We identified a total of 30 ICEs, of which 17 lack direct repeats but satisfy other criteria for ICE identification (presence of T4SS gene clusters, a *mob* relaxase and integrase sequence). Some genomes have *mob* relaxase genes flanked by direct repeats but lack an adjacent T4SS: these may be degenerate ICEs or integrative mobilizable elements (IMEs). Each genome typically carries a single ICE integrated into one of three preferred chromosomal integration sites, though UW386 and RS10 carry two ICEs. All ICEs identified in complete RSSC genomes are located on the main chromosome, none were found on the megaplasmid. Most ICEs were identified in *R. pseudosolanacearum* (23/30 ICEs). Only two were found in in R. syzygii, and five in *R. solanacearum*. All five ICEs in *R. solanacearum* are from Martinique isolates: four Rso IIB-4NPB and one Rso IIA isolate (RUN1155, Fig. 3). Strikingly, the Rso IIB-4NPB and Rso IIA ICEs are identical to each other and homologous to a 45.9kb ICE present in eight Rps genomes that collectively span the diversity of *R. pseudosolanacearum*. Since this ICE is broadly distributed among Rps genomes and also present in the reference strain Rps I-18 GMI1000, we named it ICE^RpsGMI1000^. ICE^GMI1000^ has two cargo regions of accessory gene integration we call ‘cargo-1’ and ‘cargo-2’, according to their integration site within the ICE (Fig. 4, Fig. S12).

**Fig. 4.**
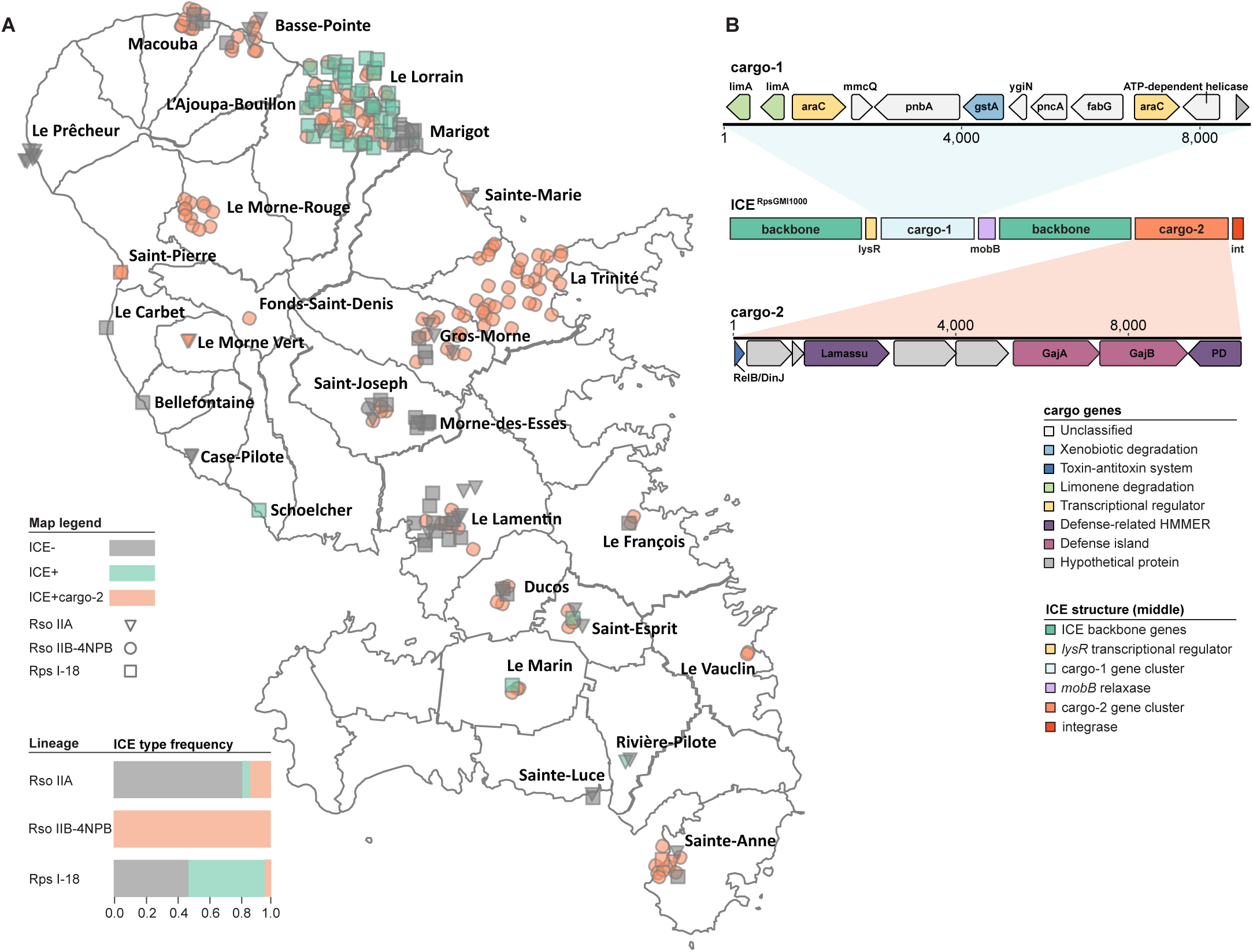
*Ralstonia* ICE distribution and cargo gene carriage in Martinique. (A) Sampling location of all Martinique isolates sequenced in this study. The exact longitude and latitute of the sampling locations were jittered for visualisation purposes. Rso IIA (triangle), Rso IIB-4NPB (circle), Rps I-18 (square) are represented with grey (ICEless), green (ICE ^GMI1000^ with conserved cargo-1 and different cargo-2) or orange (ICE^GMI1000^ with conserved cargo-1 and cargo-2) icons. The freuquency of strains carrying each ICE type is shown inset. (B) A diagram of ICE^GMI1000^ structure is shown in the top right, delineating the position of backbone and cargo gene clusters. The coding sequences are shown above (cargo-1) and below (cargo-2) for the conserved ICE, coloured according to KEGG annotation. The first backbone region includes the T4SS and has a *lysR* transcriptional regulator at the 3’ end, followed by cargo-1. The second backbone region includes the mob relaxase, *traF*, *parA*, *repA*, *qseC* and *rdfS*, along with hypotheticals and helix-turn-helix and various DUF domain-containing proteins.

We assessed the spatial structure of ICE carriage among the entire set of 7 complete and 275 draft genomes representing the densely sampled Martinique population and found local hotspots of ICE exchange in Martinique (Fig. 4). While there are multiple regions in which both lineages co-occur, it is rare for these divergent lineages to share identical ICEs (Fig. S13). At a global scale, ICE carriage appears more common among *R. pseudosolanacearum* than *R. solanacearum*, where we have only found them in lineages circulating in Martinique: Rso IIB-4NPB and Rso IIA. ICEs can be gained and lost quite rapidly however, and intensive sampling and sequencing of each lineage in Martinique shows that while ICE carriage is nearly ubiquitous among Rso IIB-4NPB, Rps I-18 carriage is variable. There is a striking hotspot of lineage and ICE co-occurrence in Le Lorrain: Rps I-18 and Rso IIB-4NPB isolated from the soil in this region carry ICEs identical in their backbone and cargo-1 genes, but variable in the cargo-2 genes (Fig. 4, Fig. S13, Fig. S14). There are additional instances of identical ICE^RpsGMI1000^ carriage in Rps I-18 and Rso IIA in Martinique, but none on the mainland. The presence of spatial hotspots of horizontal gene transfer in the environment, where mobile element exchange appears more frequently between co-occurring lineages, is mirrored by the presence of recombination hotspots within the mobile elements themselves. While cargo gene clusters vary between different ICEs, their overall function does not: cargo-1 gene clusters always encode metabolism-associated genes and cargo-2 gene clusters always encode bacterial defence systems (Fig. S14).

The metabolic cargo-1 content is identical in both Rps I-18 and Rso IIB-4NPB ICEs (e.g. RUN1145, RUN0777 and RUN1960 includes two *limA* genes encoding limonene 1,2-epoxide hydrolases^48^ (OFBKGH_09550, OFBKGH_09555 in RUN1145); a phthalate ester hydrolase *pnbA*; an antibiotic biosynthesis monooxygenase *ygiN* (also known as ABM, OFBKGH_09580); a glutathione S-transferase *gstA* linked with the detoxification of organochlorine compounds (1-chloro-2,4-dinitrobenzene) used as an herbicide and pesticides; a pyrazinamidase/nicotinamidase *pncA*, which converts nicotinamide to nicotinic acid, and *fabG*, which encodes a reducing agent of ethyl 4-chloroacetoacetate, an intermediate used in herbicide and pesticide production. Rps I-18 and Rso IIB-4NPB ICEs carry distinct cargo-2 clusters encoding multiple defence systems, such as anti-plasmid/phage Lamassu and anti-viral Gabija islands^49,50^. The only other ICE present in a Rso genome (Rso IIA RUN1155, also from Martinique) shares a similar cargo-2 region with Rso IIB-4NPB ICE^GMI1000^ (Fig. 3, Fig. S14).

### Surge of prophage acquisition in the emergent lineage

We found Rso IIB-4NPB acquired three *Caudoviricetes* prophages, one of which is uniquely present in Rso IIB-4NPB and no other sequenced RSSC genome (Table S4, Table S7). The two other prophages are present in other Rso circulating in Central and South America. All complete Rso IIB-4NPB genomes (RUN0777, RUN0823, RUN0808, RUN1145 and basal RUN2004) have the three intact (and one partial) prophage insertion on the chromosome, while the four complete Rso IIB-4 (UW163, Po82, IBSBF1503 and CIAT_078) have at most a single intact prophage on the chromosome and megaplasmid each. Panaroo pangenome exploration and Pyseer kmer-based association tests alike uncover the Rso IIB-4NPB-unique prophage and a downstream auxiliary region (Fig. S15, Table S9). Whole genome alignment of Rso IIB-4 UW163 from *Musa acuminata* x *balbisiana* AAB (Peru, 1967) and Rso IIB-4NPB RUN1145 from *Cucurbita moschata* (Martinique, 2002) confirms horizontal gene transfer in the form of prophage and ICE movement has had a significant impact on Rso IIB-4NPB genome evolution. Despite being isolated 45 years apart from different hosts and regions, these genomes are highly conserved across the chromosome and megaplasmid (Fig. S15, Fig. S16). The most striking difference between the two lineages are large insertions in Rso IIB-4NPB, most of which are predicted to be prophage. RECOPHY indicates the region with the highest SNP density in the Rso IIB-4 core genome alignment (1,060 out of 2,894 SNPs) is adjacent to two prophages and a proximal *fhaC-fhaB* locus (Rso IIB-4NPB RUN1145 chr1: 227,422-304,634). This region also contains a high density of Martinique-associated kmers in RUN1145 and *Musa*-associated kmers in UW163. The closest BLASTn hits to the novel prophage are soil isolates from Asia (*Ralstonia* metagenome-assembled genome from estuary sediment in China, *Pandoraea pnomenusa* from sludge in Malaysia and *Massilia putida* from soil in China) and *Xanthomonas albilineans*, a bacterial plant pathogen causing lethal leaf scald in sugarcane from Brazil (Fig. S17). In addition to the core genes encoding the phage head, tail, baseplate and restriction endonucleases, the Rso IIB-4NPB-specific prophage carries auxiliary genes whose products are predicted to function as a circadian clock protein (*kaiA*, OFBKGH_12090 in RUN1145) and a methyl-accepting chemotaxis protein (OFBKGH_12085). Cumulatively, the data shows ICE acquisition and prophage infections occurred multiple times in the ancestor of the emergent Rso IIB-4NPB lineage, either prior or soon after its dissemination to Martinique.

### CRISPR array reveals a history of viral encounter in the emergent Rso IIB-4NPB lineage

Evidence of repeated mobile element acquisition in the host-range expanded lineage Rso IIB-4NPB attests to the high frequency of encounter and exchange with members of the soil, freshwater and plant microbiomes. The carriage of antiviral defense gene clusters on the ICE and presence of type I-E CRISPR-Cas systems among RSSC genomes indicates the plant pathogen is frequently challenged by viral attack. CRISPR-Cas systems confer immunity to phage via the acquisition of spacer sequences from prior infections. We found that both Rso IIB-4 and Rso IIB-4NPB share conserved CRISPR-Cas operons (*cas3*, *casA/cas8e*, *casB/cas11/cse2*, *cas7e*, *cas5e*, *cas6e*, *cas1e*, *cas2*), but vary in their CRISPR arrays. The Rso IIB-4NPB CRISPR 1 and CRISPR 2 arrays are nearly ten times longer than the Rso IIB-4 arrays: Rso IIB-4 UW163 has 20 spacers compared to the 171 spacers in Rso IIB-4NPB RUN1145 (of which 11 are duplicates with 1-5bp differences). These genomes share one leader-proximal and 15 leader-distal spacers. The kmer-based association tests show that the largest cluster of Martinique-associated kmers that do not map to ICEs or prophage actually map to the leader-proximal position of CRISPR arrays in Rso IIB-4NPB, corresponding to more recent spacer acquisition than the leader-distal spacers shared with Rso IIB-4 (Fig. S15, Fig. S18)^51,52^. This represents a rapid expansion of CRISPR-mediated immunity in the emergent lineage, indicating the emerging lineage encountered phage during its expansion across Martinique.

We inferred the identity of the viruses challenging Rso IIB-4NPB using spacer sequences as queries for INPHARED phage database searches. A total of nine phage were identified using this approach (Table S8). Three phage are targeted by spacers in Rso IIB-4 and Rso IIB-4NPB alike, and six are exclusively targeted by Rso IIB-4NPB. Interestingly, five of the six phage targeted by Rso IIB-4NPB are known to infect Rps phylotype I strains (RPZH3, RS603, RSMSuper, Eline, Raharianne)^53–55^. Rso IIB-4NPB immunity to phage infecting Rps suggests they not only co-occur, but also encounter the same viral threats.

## Discussion

Outbreaks of plant disease tend not to be well sampled outside the focal host, or even deeply sampled within the focal host of interest, limiting our ability to infer outbreak origins or identify the genomic changes associated with pathogen emergence. Our knowledge of pathogen host range and the selective environments they are exposed to outside the acute infection stage is extremely limited. Infection assays using a broad panel of potential hosts provide an indication of potential host range, yet these experiments typically include high inoculum densities, mechanical damage and/or other methods of bypassing the initial invasion stage. These results are therefore not likely to predict what is observed during natural field infections. The densely sampled collection from Martinique and French Guiana is therefore unique as it captures a disease outbreak that affected a wide range of hosts, including isolates from asymptomatic and environmental reservoirs alike, across island and mainland populations in the Caribbean and South America. This work provides unprecedented insight into the evolutionary history and processes acting on a pathogen community during an outbreak event, revealing how mobile elements exchange and interactions with viral predators have shaped their genomes.

The Martinique outbreak was caused by the parallel expansion of two distantly related pathogen lineages: Rso IIB-4NPB and Rps I-18. While at present considered an unusual event^56^, this is likely to change with increased outbreak sampling and sequencing. Genetic exchange between endemic and introduced epidemic pathogens has been documented^57,3^. Interestingly, the broad host range Rso IIB-4NPB lineage emerged from a diverse endemic population that has been circulating in the Americas for some time, while the lineage that ultimately originated in Asia retains a signature of specialization on Solanaceae. Rso IIB-4NPB is currently unique in causing natural infections in the most diverse group of hosts ever documented for a single lineage in the field. Our results show this emergence event was not driven by T3E innovation and illustrates the risk of using limited sampling and whole genome sequencing to inform candidate gene selection for studies of virulence gene function. For example, a strain IBSBF1503 previously identified as Rso IIB-4NPB is in fact distantly related to the emergent lineage that swept across Martinique^58^, and there is very little variation in effector carriage between Rso IIB-4NPB and Rso IIB-4: a single hypothetical effector (*T3E_hyp3*) was acquired and two (*ripAX2* and *ripP3*) were lost. It is possible that regulatory changes or allelic variation in T3Es may alter apparent host range^59^, but the observed differences appear unlikely to account for the apparent expansion in host range compared to Rso IIB-4, particularly since single effector deletions typically have limited consequences for pathogenicity^60^. Although T3E changes associated with host range expansion in the emergent lineage may have occurred prior to its dissemination to Martinique, the ancestral mainland population from which Rso IIB4-NPB arose remains associated with *Solanaceae* infections. In sum, there is no evidence the outbreak of Rso IIB-4NPB in Martinique is driven by T3E-mediated changes altering pathogen interactions with a panoply of distantly related hosts. The genomic evidence instead shows the acquisition of anti-viral defences and novel metabolic functions are more likely to have played an important role in the successful establishment of the pathogen across a broad range of hosts during this outbreak. This does not preclude the possibility outbreaks caused by related lineages are linked with T3E acquisition, as may be the case for banana-pathogenic Rso IIB-4.

Resolving fine-scale geographic patterns of mobile element carriage revealed hotspots of ICE carriage and interspecies exchange in Martinique, and provides insight into the dynamic evolution of the mobile elements themselves. These ICEs exhibit conservation of function in both hotspots of accessory gene acquisition: despite variable gene content the cargo-1 hotspot always consists of metabolism-associated genes and cargo-2 always consists of defense-related genes. Investigation of mobile element carriage in all complete genomes available for the species complex suggests that despite chromosomal homology at preferred ICE integration sites and the presence of direct repeats targeted by the ICE^RpsGMI1000^ integrase, ICE carriage is less common in Rso and Rsy compared to Rps. Expanded population level sequencing cautions against inferences drawn from limited sequencing however, since we show MGE carriage can vary from one sampling site to another even within a single lineage. Similarly, rapid movement of ICEs prevent reliable inference of the source and directionality of transfer events, but their absence from Rso IIB-4 and near ubiquity among Rso IIB-4NPB isolates suggests ICE^RpsGMI1000^ acquisition is linked with the outbreak in Martinique. Since ICEs mediate the horizontal transfer of entire functional genetic modules, allowing rapid adaptation to novel hosts and environmental niches, the acquisition of an ICE with novel metabolic enzymes may have been pivotal to the expansion of Rso IIB-4NPB.

All Rso IIB-4NPB strains in Martinique carry an ICE with cargo-1 genes whose products are linked with the degradation of environmental pollutants, reactive xenobiotics and plant-derived metabolites. The phthalate ester hydrolase (*pnbA*) is linked with the degradation of phthalate esters, common synthetic organic compounds that are known to be environmental pollutants^61^. Homologs of *gstA* and *fabG* are known to degrade organochlorine compounds used as primary active ingredients (1-chloro-2,4-dinitrobenzene) or intermediates (ethyl 4-chloroacetoacetate) in the production of herbicides and pesticides^62,63^. Multiple cargo-1 genes have products linked with the manipulation of host-derived metabolites: the limonene 1,2-epoxide hydrolases *limA* can act on host plant epoxides, induced during both pathogen attack and methyl jasmonate application^64^; a homolog of *ygiN* (ABM) is known to hydroxylate jasmonic acid in rice, attenuating plant defences during infection^65^, and pyrazinamidase/nicotinamidase (*pncA*) is known to convert nicotinamide to nicotinic acid. The accumulation of nicotinic acid during grape xylem colonisation by *Xylella fastidiosa* suggests *pncA* may be linked with vascular pathogenesis^66^. *Ralstonia* spp. degradation of plant metabolites has elsewhere been observed in Rso IIB-1 UY031, which degrades phenolic compounds in xylem sap^67^, and many *Ralstonia* spp. genomes contain salicylic acid degradation pathways^45,68,69^. Bacterial metabolism and manipulation of nutrient availability in host plants is increasingly recognised as an important component of pathogenicity, in mammalian and plant pathogens alike^70–73^. For instance, genes whose products facilitate the use of xylem sap as a carbon source^74^ and horizontally acquired components of the denitrification pathway^75^ have been shown to be important for xylem colonisation and virulence in *Ralstonia*. Transcriptome sequencing of experimentally evolved Rps I-18 GMI1000 showed fatty acid metabolism is upregulated in evolved clones of Rps I-18 GMI1000, highlighting the importance of metabolism during adaptation to a novel host^76^. Metabolism is likely carefully regulated according to infection stage, as was shown for Rso IIB-1 UW551, which downregulates carbohydrate metabolism and tricarboxylic acid-related pathways during late stages of xylem infection^77^. The acquisition of novel pathways for substrate catabolism may enhance bacterial growth in the soil and/or within plant tissues, resulting in an explosive increase in symptom production across a wide range of hosts.

Recombination events introducing metabolic enzymes occurred beyond the ICE as well, in the ancestor of Rso IIB-4 and Rso IIB-4NPB. These are similarly linked with the degradation of either plant-derived compounds or protection against environmental toxicity or oxidative stress. The acquisition of a TauD/TfdA family dioxygenase linked with synthetic auxin degradation may confer a fitness advantage in niches frequently exposed to phenoxy herbicides. The ability to metabolise toxic or antimicrobial compounds is not unusual in *Ralstonia* and close relatives: *R. pickettii* degrades DDT pesticides ^1^ and *Cupriavidus necator* (formerly *R. eutropha*) degrades chloroaromatic compounds^78^. The ancestral acquisition of a carboxymuconolactone decarboxylase is also of considerable interest, since the protocatechuate catabolism pathway is responsible for the conversion of lignin-derived aromatic compounds into amino acids and organic acids, potentially fueling *Ralstonia* growth in plant tissues.

Emerging pathogens may themselves serve as hosts for potential parasites. The antiphage defense in ICE cargo-2, expansion of CRISPR repeats and acquisition of multiple fragmented prophage in Rso IIB-4NPB attests to frequent viral encounter, and indicates immunity to predatory phage is likely to provide an advantage to the pathogen^79^. Phage resistance is similarly implicated in the success of 7^th^ pandemic *Vibrio cholerae* lineages^80,4^. Since some of the Rso IIB-4NPB CRISPR spacers with identifiable origins correspond to phage known to successfully infect *R. pseudosolanacearum* strains like Rps I-18, coinfections are likely sites of exposure to lytic phage, along with the acquisition of ICEs and prophage carrying accessory genes altering microbial lifestyle and pathogenicity^81^. Our findings emphasize the value of integrating genomic, ecological and evolutionary perspectives to understand disease emergence. As agricultural intensification continues to reshape plant-microbe interactions, pathogen lineages that can rapidly acquire and integrate novel genetic material may be more likely to exploit novel niches and hosts. This work underscores the need for field surveillance strategies that extend beyond focal pathogen populations to the broader community and reservoir of genetic diversity that fuels their evolution.

## Methods

### Genome sequencing

*Ralstonia* spp. were collected in Martinique and French Guiana as described in Wicker *et al*. 2009 and Deberdt *et al*. 2014. Isolates collected after the initial sampling were assigned to phylotype and sequevar by sequencing endoglucanase (*egl*) and mutation mismatch repair (*mutS*) genes. For this study, we extracted genomic DNA from 487 strains using Promega Wizard, then used a modified Nextera protocol for library preparation^82^. Paired-end sequencing with Illumina NextSeq 2000 was then performed. Since the first round of sequencing produced under 25-fold average coverage and under 40-fold maximum coverage, a second round of library preparation and sequencing was performed. Eight isolates (RUN0777, RUN0808, RUN0820, RUN0823, RUN1145, RUN1155, RUN1952 and RUN2004) were additionally sequenced using Oxford Nanopore Technology with a FLO-MIN106 flow cell and SQK-RBK004 barcoding kit. Basecalling and demultiplexing was performed with guppy_basecaller (v. 5.0.7) and guppy_barcoder v. 4.5.4^83^.

### Genome assembly and annotation

Paired-end 150bp reads of both Illumina sequencing runs were filtered and trimmed with fastq_screen^84^, bbduk^85^ and fastp^86,87^ (--dedup --detect_adapter_for_pe --cut_front 15 --length_required 50). Assembly was performed with SPAdes^88,89^ using both separate and pooled (i.e. both sequencing runs) read sets. QUAST was used for assembly quality assessment^90^. Pooling both sequencing runs improved average assembly N50 values from 73Kb to 195Kb and decreased average contigs per assembly from 370 to 217, so assemblies generated from pooled sequencing runs were used. After filtering out assemblies with under 50-fold coverage, over 250 contigs, total assembly length exceeding 6Mb and N50 exceeding 75Kb, a total of 409 assemblies were retained for downstream analyses.

Oxford Nanopore long read data was filtered with Filtlong (v. 0.2.1)^91^ to remove reads under 1Kb and the 5% of reads with the lowest quality values, then assembled with Trycycler (v. 0.5.4)^92^, using Flye^93^, Miniasm^94^ with Minipolish^95^ and Raven^96^ as underlying assemblers. Contigs that did not pass the reconciling step of Trycycler pipeline were manually removed. Final Trycycler consensus sequences were polished three times using long reads with Medaka^97^, followed by two rounds of short read polishing using BWA^98,99^ and Polypolish^95^, using separate batches of the Illumina reads produced by each sequencing run.

All assemblies were annotated by Bakta^100^ using Prodigal^101^ gene models and a reference protein database built using all complete *Ralstonia* spp. genome assemblies available from the RefSeq database in October 2022.

### NCBI datasets and reference genomes

All available *R. pseudosolanacearum, R. syzygii* and *R. solanacearum* genome assemblies were downloaded from NCBI in February 2023 to provide additional phylogenetic context for analyses. Assemblies with fewer than 200 contigs, total assembly length between 5.0 - 6.2Mb, N50 over 75Kb and unflagged by the NCBI Genome^102^ for evidence of potential contamination were retained. A total of 410 *Ralstonia* spp. genomes passed filter (Table S1). Three isolates were present in duplicate in the NCBI filtered dataset (MAFF 211479, UW163 and UW551), the lowest quality duplicate assembly was removed, producing a final NCBI dataset of 407 assemblies. These were then reannotated with Bakta and merged with the 409 assemblies produced in this work for downstream pangenome analyses. Metadata (host, geographical region and year of isolation) for genomes in the NCBI dataset was obtained by inspecting Biosample entries. If the information was missing in Biosample entries, we used RSSCdb records^103,29^ or publications describing the initial isolation of the strain of interest. Metadata for the strains CIP128_UW455, UW258 and NCPPB282_UW491 was not used for the visualization due to discrepancies.

### *Solanaceae* specialisation test

Level of specialisation toward *Solanaceae* family hosts among three *Ralstonia* groups (Rso IIA, Rso IIB-4NPB and Rps I-18) on Martinique was estimated by two Chi-squared Test for Independence using function chi2_contingency from scipy package. Contingency table of infection frequencies was composed from numbers of Rso IIA, Rso IIB-4NPB and Rps I-18 isolates from *Solanaceae* and non-*Solanaceae* plants sampled on Martinique. Environmental isolates and the isolates with missing metadata were not included. Tests were performed for two pairs Rso IIB-4NPB vs Rps I-18 and Rso IIB-4NPB vs Rso IIA with the Null hypothesis “Probability of infecting *Solanaceae* family plan is independent from the *Ralstonia* lineage”. Observed and expected distribution were visualized by matplotlib package.

### Type 3 secreted effector identification

Amino acid sequences of 118 *R. solanacearum* type 3 secreted effectors (T3Es) from the manually curated RalstoT3E database^104^ were downloaded and used as query sequences for DIAMOND BLASTp^105^ searches against proteomes of the Bakta-annotated assemblies, using the --ultrasensitive mode. Hits were filtered using a S30L25 strategy: retaining sequences with alignment identity of at least 30% and query sequence coverage of 25% or greater. Since the T3E query set includes multiple alleles of the same effector families, hits were then grouped into T3E families by sorting first by identity and query length coverage. T3E family assignment for each protein sequence was performed using the most significant T3E hit. Effector presence and absence was visualised by a heatmap in R, omitting duplication events. This approach was tested by comparing the predicted T3E repertoires in Rps I-18 GMI1000 and Rsy RPSI07 with the ones provided in the RalstoT3E database. Average pairwise similarity of T3E repertoires within RSSC clades was estimated by Scikit-learn function cosine_similarity^106^ using the information about presence-absence of effector families as an input binary matrix.

### Pangenome inference and core gene tree reconstruction

Panaroo^107^ was used to identify core and flexible genomes for the RSSC dataset, i.e. the merged dataset of 816 *Ralstonia* genomes sequenced in this study (409 strains) and downloaded from NCBI (407 strains). A concatenation of core genes was used as an input alignment for phylogenetic analysis of the species complex. Alignment columns with gaps or ambiguities and invariant sites were removed with SNPsites^108^, producing a final alignment of 137,423bp. Tree inference was then performed using IQtree^109^ with a GTR+ASC model of sequence evolution, model for alignments that do not contain constant sites, and 1000 bootstrap replicates. The ggtree R package^110^ was used for all tree and metadata visualisation.

Panaroo gene presence and absence tables for the RSSC dataset were queried using a custom Python script to identify genes uniquely present in either Rps I-18, Rso IIB-4 or Rso IIB-4NPB. Genes are referred to as clade-specific when they are present in at least 95% of all genomes in a given focal clade, and present in no more than 10 non-focal clade assemblies.

The species complex tree from the set of core genes identified by Panaroo was used for clade delineation and lineages assignment. Most genomes clustered according to the phylotypes initially assigned by *egl* and *mutS* sequencing in Wicker *et al.* 2007 and Deberdt *et al.* 2014, although some discrepancies are present. We found previous sequevar designations can provide an inaccurate view of strain relatedness, e.g. RUN2182 is classified as a Rps I-48, but groups with Rps I-47 in the core gene tree. Some strains were also reclassified into different phylotypes based on the core gene tree and RUN1259 was initially classified as Rso IIB but is in fact Rso IIA (Fig S1., Table S1). The clade designation in this work may therefore differ from historically assigned sequevar or even phylotype labels, since these were first assigned using *egl* and *mutS* sequences rather than whole genome sequencing and core gene tree reconstruction. The criteria for membership in a named clade is exceeding a minimum shared identity of 99% in the RSSC core gene alignment.

Separate pangenome analyses and core gene tree reconstructions were performed on three specific clades: *R. pseudosolanacearum* phylotype 1 sequevar 18 (Rps I-18, 102 strains), *R. solanacearum* phylotype 2 sequevar 4 (Rso IIB-4, 226 strains), and the emergent lineage *R. solanacearum* phylotype 2 sequevar 4 Non-Pathogenic on Banana (Rso IIB-4NPB, 180 strains). The strains forming each clade were identified with reference to the RSSC core gene tree (Figure 1). Assemblies for IBSBF_2571 (GCA_003590605.1) and O12BS_UW407 (GCA_023076615.1) were excluded due to abnormally long terminal branch lengths and P673 was excluded due to contradictory phylogenetic position of two assembly versions: the most recent P673 assembly GCA_015698365.1 was assigned to Rso IIB-4NPB clade, while the earlier version GCA_000525615.1 grouped with other Rso IIB-4 *Epipremnum* isolates from the USA.

An additional iteration of clade-specific pangenome analysis was performed for both Rso IIB-4NPB and Rps I-18. Although strain IBSBF1503 (GCA_001587155.1) groups with other Rso IIB-4NPB strains in the RSSC core gene tree, the clade-specific tree revealed this strain is divergent and more properly referred to as an outgroup strain, therefore this was excluded from a second iteration. Similarly, Rps I-18 strains CIP266_UW505 (GCA_023075875.1) and UW393 (GCA_023076675.1) form a distant outgroup to other Rps I-18 strains, therefore these were excluded from the second iteration of the Rps I-18 pangenome analysis.

### Mobile element prediction in reference genome assemblies

All potential mobile elements, including genes annotated as integrases, transposases, prophage-related genes and IS elements, were identified and masked for the downstream phylogenetic analysis in the Bakta-annotated genome assemblies for Rps I-18 strain GMI1000 (GCA_000009125.1), Rso IIB-4 strain UW163 (GCA_001587135.1) and Rso IIB-4NPB strain RUN1145. Prophage and IS elements were predicted using the PHASTER, PHASTEST^111–113^ and ISfinder^114^ online tools. All gene annotations containing keywords “phage” OR “transposase” OR “tail” OR “head” OR “integrase” OR “terminase” OR “conjugal” OR “integrase” OR “integrative conjugative element” OR “conjugative transfer” OR “conjugative coupling” were manually searched for and flagged in Geneious Prime^115^. The results were converted into BED files, then sorted, merged and used for masking with Bedtools^116^. Masked references were used for reference-based contig mapping.

### Phylogenetic analyses

Reference-based contig mapping was performed for the Rps I-18, Rso IIB-4 and Rso IIB-4NPB clades, as defined by using RSSC and clade-specific core gene trees. Rps I-18 included 100 genomes, with GMI1000 as reference. The Rso IIB-4 clade contained 226 genomes, with UW163 as reference. Rso IIB-4NPB included 179 genomes, with RUN1145 as reference. The Brazilian *Cucumis sativus* isolate IBSBF1503 is referred to as Rso IIB-4NPB in the literature and initially grouped with Rso IIB-4NPB in the species complex tree, and, so we initially used this strain as a reference for Rso IIB-4NPB. While more closely related to Rso IIB-4NPB than any other available complete *Rso* genome, we found IBSBF1503 is clearly an outgroup relative to other Rso IIB-4NPB isolates (Fig. S3). The average pairwise nucleotide diversity is 0.002% between Rso IIB-4NPB genomes, and an order of magnitude greater (0.02%) relative to IBSBF1503. According to progressiveMauve whole genome alignments, Rso IIB-4NPB isolate RUN1145 differs from RUN1960 and IBSBF1503 by 134 and 1,215 SNPs, respectively. Since IBSBF1503 is too divergent to be considered a member of the same lineage, we reclassified IBSBF1503 as Rso IIB-4. We resequenced five Rso IIB-4NPB strains from Martinique using Nanopore long-read sequencing to generate closed hybrid assemblies and used the complete assembly of Rso IIB-4NPB RUN1145 as a reference for more reliable core genome alignment-based phylogenetic inference of relationships between closely related strains. The Rps I-18 contig mapping using Rps I-18 GMI1000 as reference resulted in 4,760,272 bp consensus alignment with 311 SNPs and 116 parsimony informative sites.

Reads and contigs mappings were performed separately for each assembly with bwa-mem2^98,99^ or Minimap2^94^ using asm5 mode respectively. All resulting BAM alignments were sorted and indexed with Samtools^117^. BCFtools mpileup was used to filter and retain bases with mapping quality above 60^118^. Variants were called with bcftools-call and filtered by bcftools-filter using the expression TYPE=“snp” && DP>=X && QUAL>30 && AF1>=0.95 with X=10 for reads and X=1 for contig mapping. Any alignment column with a gap site was removed. Consensus sequences were obtained using bcftools-consensus, masking the sites with coverage depth below one as identified by Mosdepth^119^. The consensus sequences for all strains in a given clade were concatenated into a FASTA alignment file. Sites containing N characters were trimmed with goalign clean-sites^120^. Recombinant regions were predicted and removed by ClonalFrameML^121^ using an initial tree calculated by IQtree^109^ under GTR+F+I+R2 substitution model. The final Maximum likelihood trees were calculated from the non-recombinant alignments for each clade by IQtree using the MFP (ModelFinder Plus) option for the best model search^122^. The contig mapping approach resulted in better resolved and bootstrap supported trees, therefore the trees generated by reads mapping were excluded from further analysis. Tree visualisations were created with ggtree^110^.

### Tree dating

Root-to-tip regression and dating of the Rso IIB-4NPB and Rps I-18 non-recombinant core genome trees were performed with BactDating using 10,000 Markov Chain Monte Carlo iterations^123^. TempEst was used to confirm root-to-tip regression results obtained with BactDating, setting the ‘Best-fitting root’ option of TempEst to ‘R squared’, and for outlier detection by investigating ‘Residuals’ plots^124^. Tree tips with missing sampling date information (Rps I-18 UQRS637_UW745 and UW193, Rso IIB-4NPB RUN0283 and RUN0739) and outliers (Rps I-18 UW393, GMI1000, RUN1974, UW299, RUN1975, RUN1958, RUN1955, RUN0037, RUN0679 and CIP266_UW505, Rso IIB-4NPB RUN0003, RUN0002, RUN2227, RUN2226, RUN1959, RUN1960, RUN0016, RUN2007, RUN2009, RUN2004, RUN2011, RUN0762, RUN1046, RUN1121, RUN0005, RUN1094, RUN1064, RUN1951, RUN1968, RUN1105, RUN1167 and RUN0006) identified by TempEst were dropped from the Rps I-18 and Rso IIB-4NPB alignments and trees were re-calculated. Branches with edge lengths of zero were manually collapsed into a single node by Dendroscope^125^. A Dates Permutation Test was performed 500 times for each tree with BactDating by randomly generating dates between 2000 and 2012 and randomly assigning these to tree tips while keeping the tree root (option: updateRoot=F). The simulated trees were compared to the actual dating model using the modelcompare function of BactDating.

### CRISPR-Cas system identification

CRISPR-Cas systems were identified with Bakta annotation of Rso IIB-4 and Rso IIB-4NPB genomes. CRISPR repeats and spacers were then annotated in two closed references Rso IIB-4-UW163 and Rso IIB-4NPB-RUN1145 using the CRISPRloci workflow^126^. The spacer sequences identified with CRISPRloci were then used for BLASTn searches against the RefSeq Phage database downloaded from PhageScope^127^, NCBI Nucleotide collection (nr_nt) and Environmental samples (env_nt) databases accessed in June 2024. The extracted spacers were also used for INPHARED database^128^ searches using the SpacePHARER^129^ toolkit in August 2024.

### Inter and intraspecific recombination prediction

Potential homologous recombination events within and between Rso IIB-4 and Rps I-18 lineages were investigated using the RECOPHY toolkit^130^. Two datasets were analysed: Rso IIB-4 alone and in combination with Rps I-18, using Rso IIB-4NPB-RUN1145 as a reference in each case. Prior to RECOPHY execution, all genome pairs with a Mash distance^131^ below zero were identified and one member of each pair was then randomly removed from the dataset. The final datasets included 49 genomes for the Rso IIB-4 analysis alone, and with an additional 64 Rps I-18 genomes for the merged analysis. The regions of core alignment with increased and reduced density of SNPs per 1 kb block were extracted and investigated separately. In order to find discrepancies with the core phylogeny, a simple Neighbor-Joining tree was calculated for each of these regions using Geneious Prime software^115^.

### Pangenome-wide association testing

Scoary and Pyseer pan-GWAS analyses were used to investigate associations between gene presence and absence patterns and the host or location of strain isolation^132,133^. The broad (country) and fine-scale (village) locations and all taxonomic levels of the host of isolation or environmental substrate were converted from the metadata table into a binary traits table using the pandas.get_dummies function. For example, a strain isolated from a tomato plant in China would receive the value “1” in the columns China, *Eudicots, Asterids, Solanales, Solanaceae, Solanum* and *S. lycopersicum*, and receive the value “0” in all other columns of the traits table. This traits table was used as input for both Scoary and Pyseer pan-GWAS analyses, along with the gene presence and absence table from Panaroo.

The significance of candidates identified with Scoary was assessed after multiple test corrections: individual (naive) p-value, Bonferroni adjusted p-value, Benjamini-Hochberg adjusted p-value, entire range of pairwise comparison p-values, and empirical p-value from 100 permutations, using thresholds of p=0.05 for all corrections. Significant genes identified by Scoary after all the corrections were further filtered using 95% sensitivity and 70% specificity thresholds. A lower specificity threshold was chosen due to the nature of the trait dataset: the traits table simply reflects the known host of isolation and the location of isolation. The actual host range of a particular strain may include additional hosts, closely or distantly related, just as closely related strains may be present in vastly different geographic regions. Since the Scoary and Pyseer pan-GWAS analyses aim to identify genes linked with survival in a particular host, environment or substrate, the sensitivity should be high. Those same genes may also be present in strains isolated from other plants however, therefore specificity is reduced. For example: a gene whose product is strongly associated with tomato (*S. lycopersicum*) infection should be overrepresented among tomato isolates in the dataset. Therefore we seek genes present in 95% of tomato isolates. We reduced the specificity threshold to 70% to allow for instances where another strain carries that same gene and is also capable of infecting tomato, but happens to have been isolated from a different host.

For the Pyseer analysis, the distance matrix from the RSSC core gene tree and the number of patterns for the likelihood ratio test (LRT) were calculated using phylogeny_distance.py and count_patterns.py scripts downloaded from the Pyseer GitHub page, accessed on January 2024^133^. The pattern count resulted in 6,075 patterns for the RSSC, leading to the likelihood ratio test (LRT) p-value threshold of 3.8e-08 = 0.05/(patterns_number*traits_number). Hits with effect size (beta value) under 0.2 and phenotypic variance values (variant_h2) above 0.5 were filtered out.

Pan-GWAS of gene presence/absence data is an effective tool for the identification of genes associated with traits of interest, but is limited to the accessory genome and does not test for trait associations within the core genome and intergenic regions. This gap is filled with a kmer-based association test. Analogously to the method used for gene presence/absence data, Pyseer was applied to identify the kmers significantly associated with any host or location of isolation across the RSSC, using the same binary metadata table as described for the pan-GWAS analysis. The initial set of 31bp long kmers was generated for all the RSSC genomes using unitig-caller ^1^. The distance matrix from the species complex core genes tree and the number of patterns for the likelihood ratio test (LRT) were calculated using Pyseer phylogeny_distance.py and count_patterns.py scripts^133^. The pattern count reported 1,484,018 patterns for the RSSC kmers, producing a LRT p-value threshold of 5.0e-10 = 0.05/(patterns_number*traits_number). Hits with effect size (beta value) under 0.2 and phenotypic variance values (variant_h2) above 0.5 were filtered out. The presence and positions of significant kmers in Rso IIB-4NPB reference strain RUN1145, Rso IIB-4 reference strain UW163 and Rps I-18 reference strain GMI1000 was identified and stored in BED files using a custom Python script. For visualisation purposes, all kmers mapped on a genome within a 1kb distance were merged using the bedtools-merge function^116^. The panGWAS and kmer-based tests identify many genes associated with isolation from specific hosts. These tests vary in how they control for population structure, but some genes were predicted to have associations with particular traits using multiple methods. Separate association tests were performed using either spatial location or host/substrate of isolation metadata. We noticed that many genes have overlapping predictions using both metadata types, particularly where it concerns isolates recovered from soil or water. We also found overlapping host associations at multiple taxonomic levels, for example gene presence associated with *S. lycopersicum* isolation often overlaps with gene presence associated with *Solanaceae* isolation. All overlaps were investigated using a custom Python script and visualised in an Upset plot using the UpSetR R package^134^. For the genes assigned to multiple nested traits, the lowest level of taxonomic identification (e.g. species rather than genus) or the finest level of geographical scale (e.g. village rather than country) for which significance was identified were chosen. Overlaps within the Scoary, Pyseer pan-GWAS and kmer-based outputs were identified using the same strategy.

### Identification of integrative and conjugative elements

Complete integrative and conjugative elements were identified in 133 complete RSSC genome assemblies using the following criteria: the presence of a type 4 secretion system (T4SS) required for conjugal transfer between strains, a *mob* family relaxase and integrase, and the presence of 8bp direct repeats (TTTTTTAT) encompassing the predicted ICE^135^. This 8bp repeat is described as an ICE insertion site for Rps I-18 GMI1000 in the ICEberg 3.0 database^136^. We note this repeat occurs between 50-70 times elsewhere in each RSSC genome. This does not preclude them from being actual ICE integration sites: despite the apparent existence of preferred ICE integration sites in the RSSC, some ICE integrases are known to be promiscuous and may target different repeats as attachment sites across the genome. For example, ICE Tn916 targets AT-rich loci and integrates almost at random across genomes^137^. MOBscan was used for identification of relaxases^138^, followed by BLASTn sequence homology searches for a T4SS and adjacent integrases^139^ using Rso IIB-4NPB RUN1960 T4SS and integrase sequences as BLASTn queries, in turn followed by T4SS locus confirmation with the TXSScan module of MacSyFinder toolkit^140^. The flanking repeat sequence TTTTTTAT was identified using a custom python script. In genomes where the repeat was not identified within 10kb of the T4SS, relaxase and integrase, putative insertion sites were manually identified.

The genomic context of putative ICEs in the RSSC assemblies was inferred by aligning 50 kb upstream and downstream flanking regions surrounding putative ICEs in Rps I-18 GMI1000, Rso IIB-4NPB RUN1145 and Rso IIB-4NPB RUN1960 to the genomes. The presence of genes significantly associated with isolation from Martinique by panGWAS analysis was also confirmed by BLASTn, retaining hits with e-values under 10^-5^, query length coverage over 10% and overall alignment length exceeding 1 kb.

The involvement of ICE cargo genes in specific metabolic pathways was predicted using KEGG Orthology assignment with protein sequences from the cargo regions of interest, performing searches in the database for the genus *Ralstonia* (taxa ID: 48736). The locus structure of putative ICEs was visualised with the ggtree::geom_motif R function^110^. Complete ICE sequences identified in *Rps* 362200, *Rps* RS24, *Rps* RUN1952, Rps I-18 RUN820, Rps I-18 GMI1000, Rso IIB-4NPB RUN1960, RUN0777, Rso IIA RUN1155 and *Ralstonia pickettii* strain 12J were aligned to each other using the progressive Mauve algorithm implemented in Geneious^141^.

### Geographical metadata display

Information about the geographical region of isolation for each strain was obtained from the country of isolation metadata collected for the RSSC strains set as described above. Counties of isolation were converted into continents using pycountry Python package^142^. The “Caribbean” region of isolation was manually assigned for all Martinique isolates to highlight the region of interest.

Longitude and latitude of the sampling site was recorded for 83/500 strains sampled in Martinique and French Guiana. For the remaining 417/500 strains, the commune (administrative district) or the nearest village was listed in the strain metadata. For these cases, approximate longitude and latitude values were retrieved using the geolocator function of the geopy.geocoders Python package using the commune or nearest village names^143^. Commune maps of Martinique and French Guiana in geojson format were downloaded from the France Geojson collection^144^. The actual and estimated longitude and latitude values for each isolate were visualised with the geopandas package^145^.

### Data availability

Sequence data generated for this work is available via NCBI SRA Accession PRJNA1196686. Annotated assemblies are available via Zenodo at https://zenodo.org/records/15619409.

## Supporting information

Supplementary Results and Figures S1-S18

## Acknowledgements

We acknowledge the support of the Max Planck Society in funding the Independent Research Group on Plant Pathogen Evolution (H.C.M.) and for providing an International Max Planck Research School Ph.D. scholarship (D.E.). Research was supported in part by grant NSF PHY-2309135, the Gordon and Betty Moore Foundation Grant No. 2919.02, and the Chan Zuckerberg Initiative DAF grant to the Kavli Institute for Theoretical Physics (KITP). This work was performed in accordance with the Nagoya Protocol on Access and Benefit Sharing (ABSCH-IRCC-FR-254778-1). We thank Gal Ofir and Sheila Roitman for helpful discussion, Ludovico Calabrese for assistance with Recophy and members of the McCann lab for discussion and comments on the manuscript.

## Author contributions

D.E., Y.P., S.P., E.W. and H.C.M designed research, D.E., Y.P., M.K., C.W., C.F. and M.V.-Z. performed research; D.E., M.K., E.C., Y.F.C., E.W. and H.C.M. analysed data; D.E. and H.C.M wrote the paper; all authors commented upon and/or approved the manuscript.

