## Supplementary Results and Figures S1-S18 for "Interspecies exchange of a mobile genetic element during a plant disease outbreak"

**Interlineage transfer of a mobile genetic element during a plant disease outbreak**

### Supplementary Figures

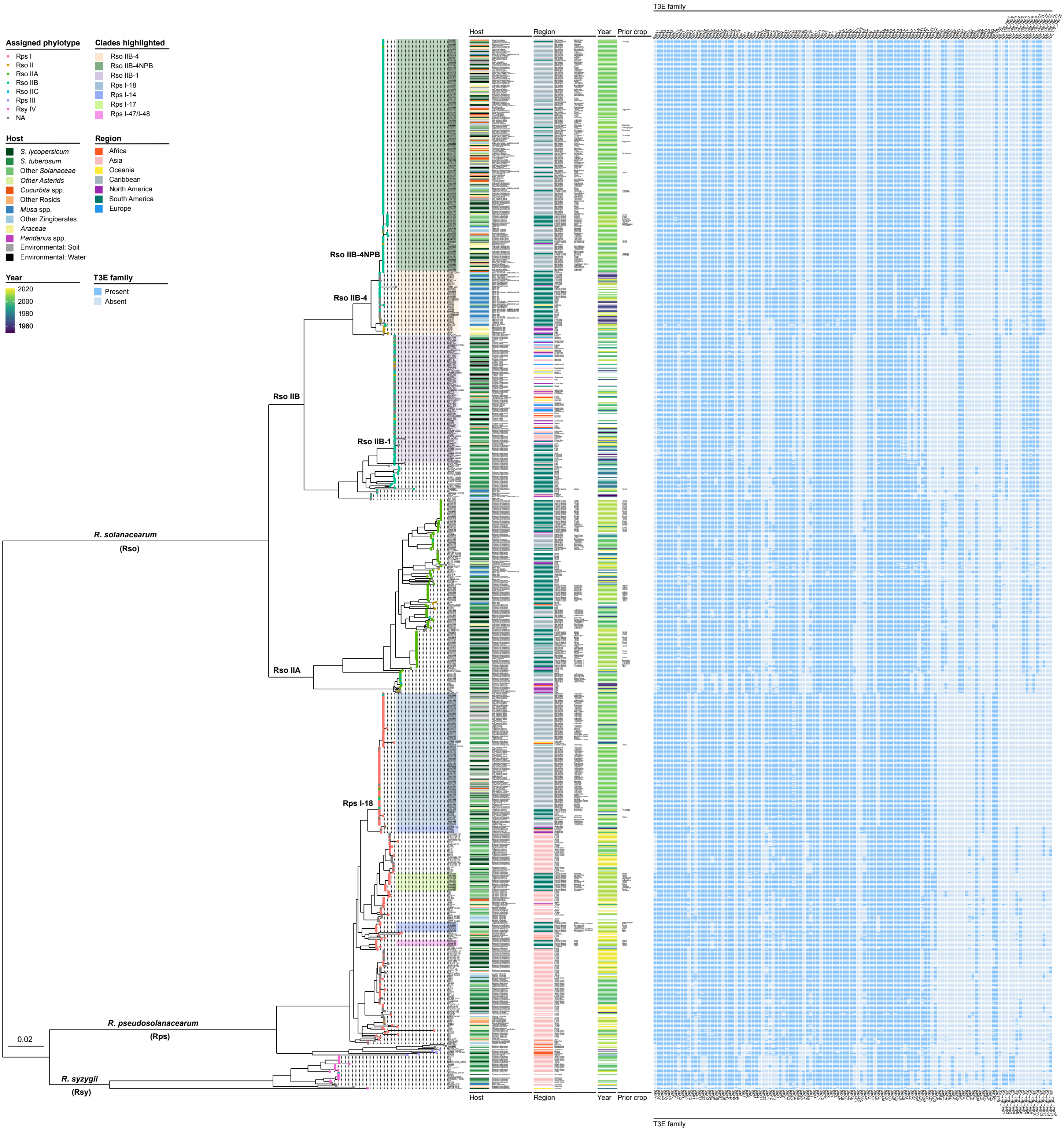

**Fig. S1. Core gene tree of the *Ralstonia solanacearum* species complex.** Maximum likelihood tree of all RSSC genomes available in this study. Nodes with bootstrap support below 70 are collapsed. Clades representing Rso and Rps lineages of interest are highlighted in the tree. Tree leaves are labelled by strain names and dots coloured according to the phylotype indicated in BioSample entry for the strains from NCBI or assigned by *egl* sequence for the strains from Martinique and French Guiana. Labels of reference strains and the eight complete or near-complete assemblies generated in this study are shown in bold. Host and geographical area of isolation are displayed together with text labels showing the species, country and commune of isolation if available. The year of isolation and common name of the crop grown in that location the preceding year (preceding crop) are shown for each strain, if known. The presence and absence of the full set of known RSSC T3E families are shown at right.

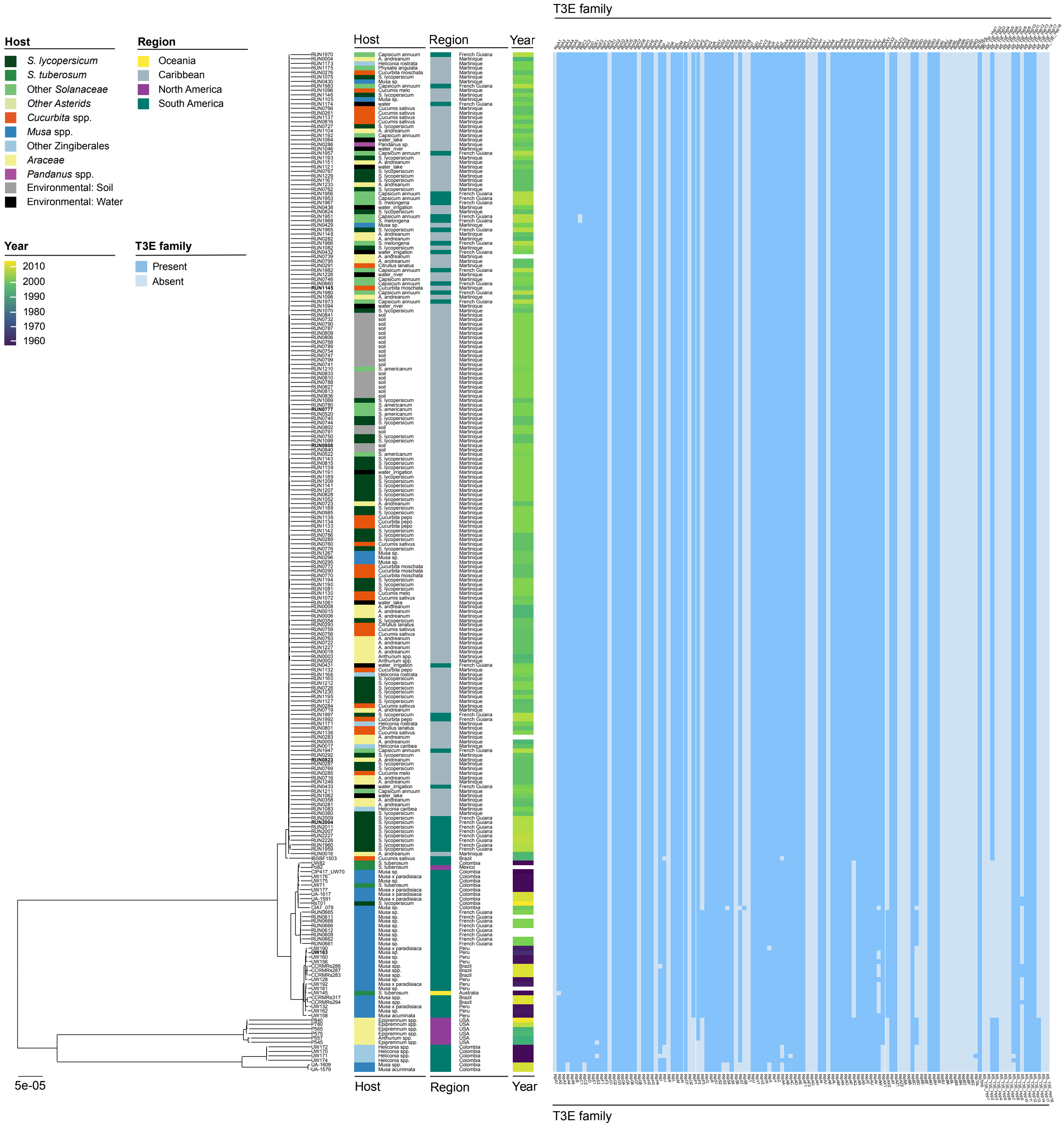

**Fig. S2. Rso IIB-4 core genome tree with T3E families.** Maximum likelihood tree of all Rso IIB-4 genomes calculated from contig alignment against UW163 as reference. Nodes with bootstrap support below 70 are collapsed. Host, geographical region and year of isolation are shown as in Fig. S1. Complete or near-complete genome assemblies generated in this study are shown with bold strain labels. Presence and absence patterns of T3E families are shown at right.

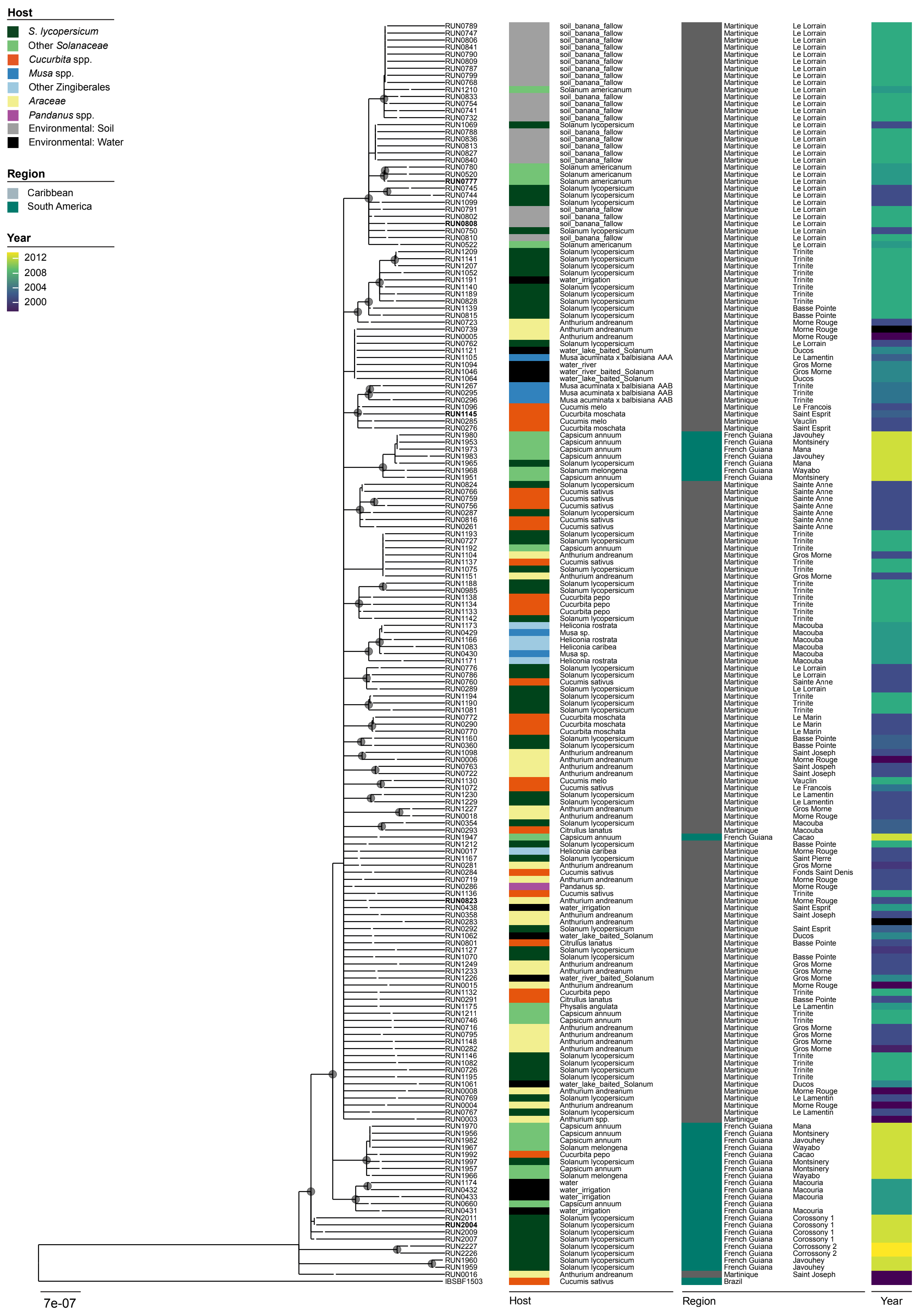

**Fig. S3. Rso IIB-4NPB core genome tree with outgroup.** Maximum likelihood core genome tree of all Rso IIB-4NPB isolates using IBSBF1503 as both reference and outgroup genome. Nodes with bootstrap support values below 70 are collapsed. Host, geographical region and year of isolation are shown as in Fig. S1.

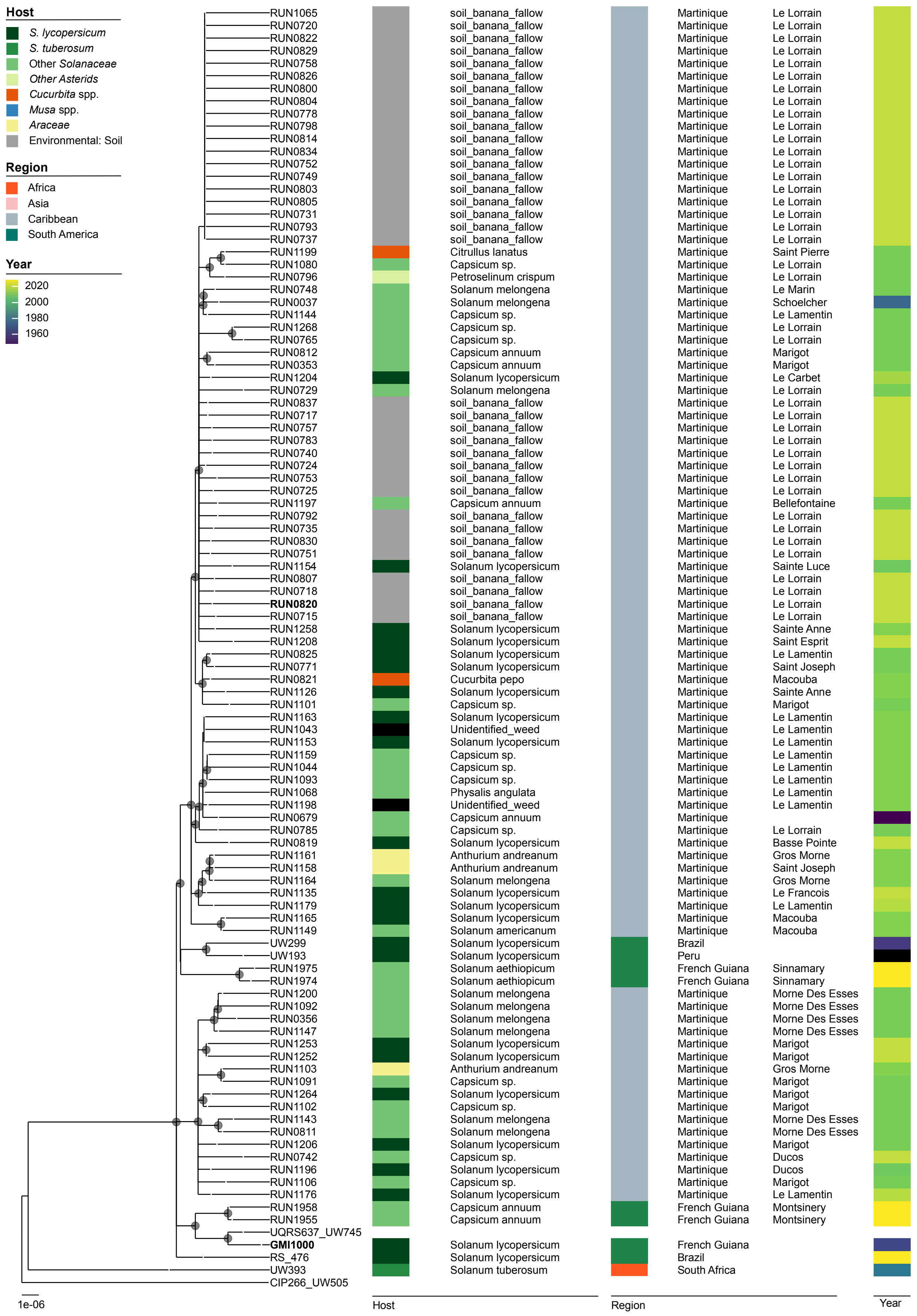

**Fig. S4. Rps I-18 core genome tree with outgroup.** Maximum likelihood core genome tree of all Rps I-18 with UW393 and CIP266_UW505 included as outgroup strains, using Rps1S18 GMI1000 as reference. Nodes with bootstrap support below 70 are collapsed. Host, geographical region and year of isolation are shown as in Fig. S1. Complete or near-complete genome assemblies generated in this study are shown in bold.

A.

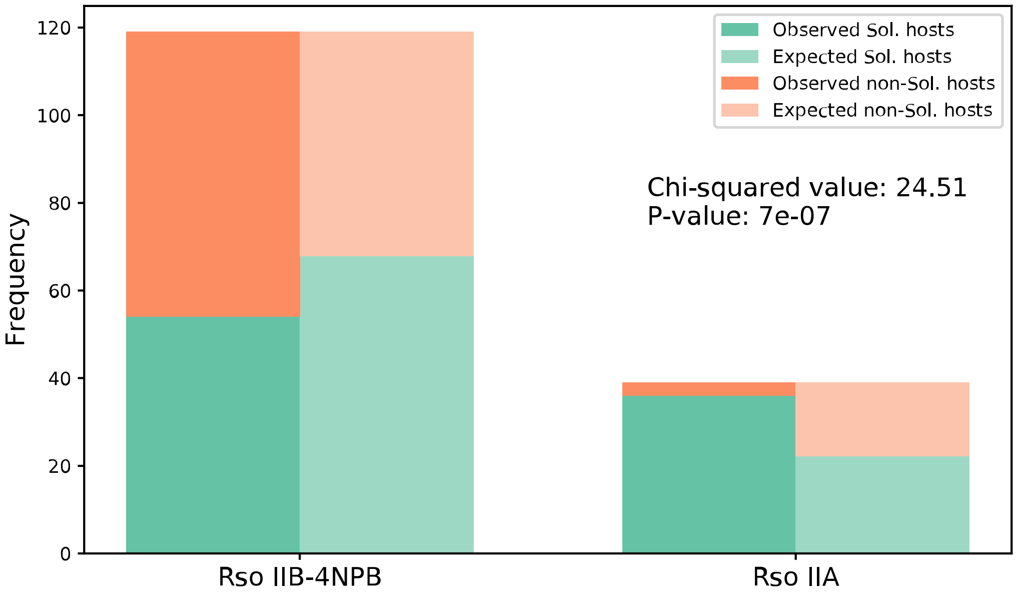

B.

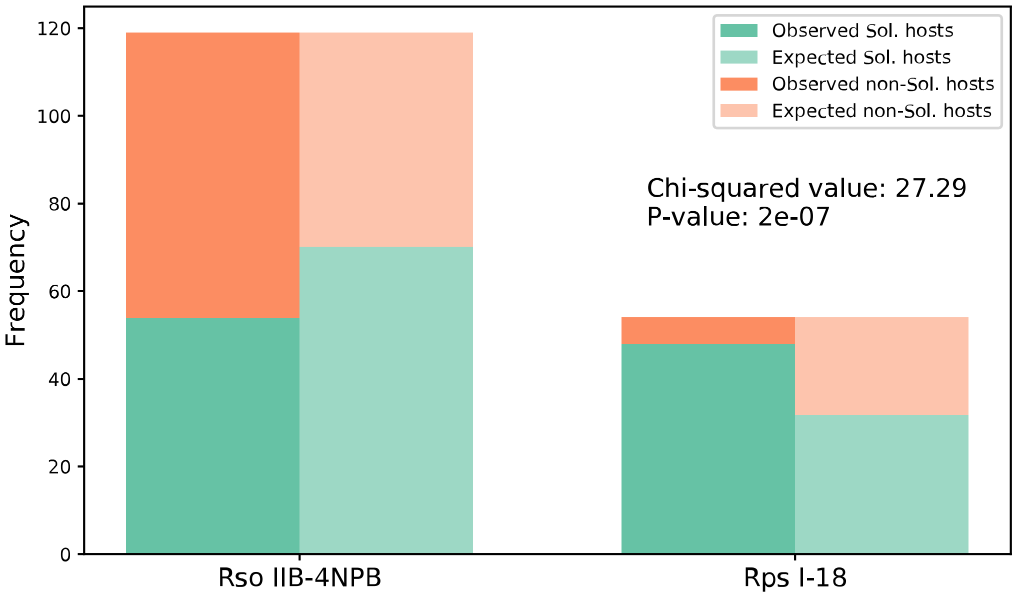

**Fig. S5. Observed vs. Expected infection frequencies by species.** Chi-squared Test for Independence comparing level of specialization toward Solanaceae family hosts between (A) Rso IIB-4NPB and Rso IIA, and (B) Rso IIB-4NPB and Rps I-18 populations on Martinique. Contingency table of infection frequencies was composed from counts of isolates from Solanaceae (green) and non-Solanaceae (orange) plants sampled on Martinique. Environmental isolates and the isolates with missing metadata were not included. Tests were performed with the Null hypothesis "Solanaceae infection probability on Martinique is independent of pathogen lineage". Actual and expected distributions shown in bold and faint colors respectively.

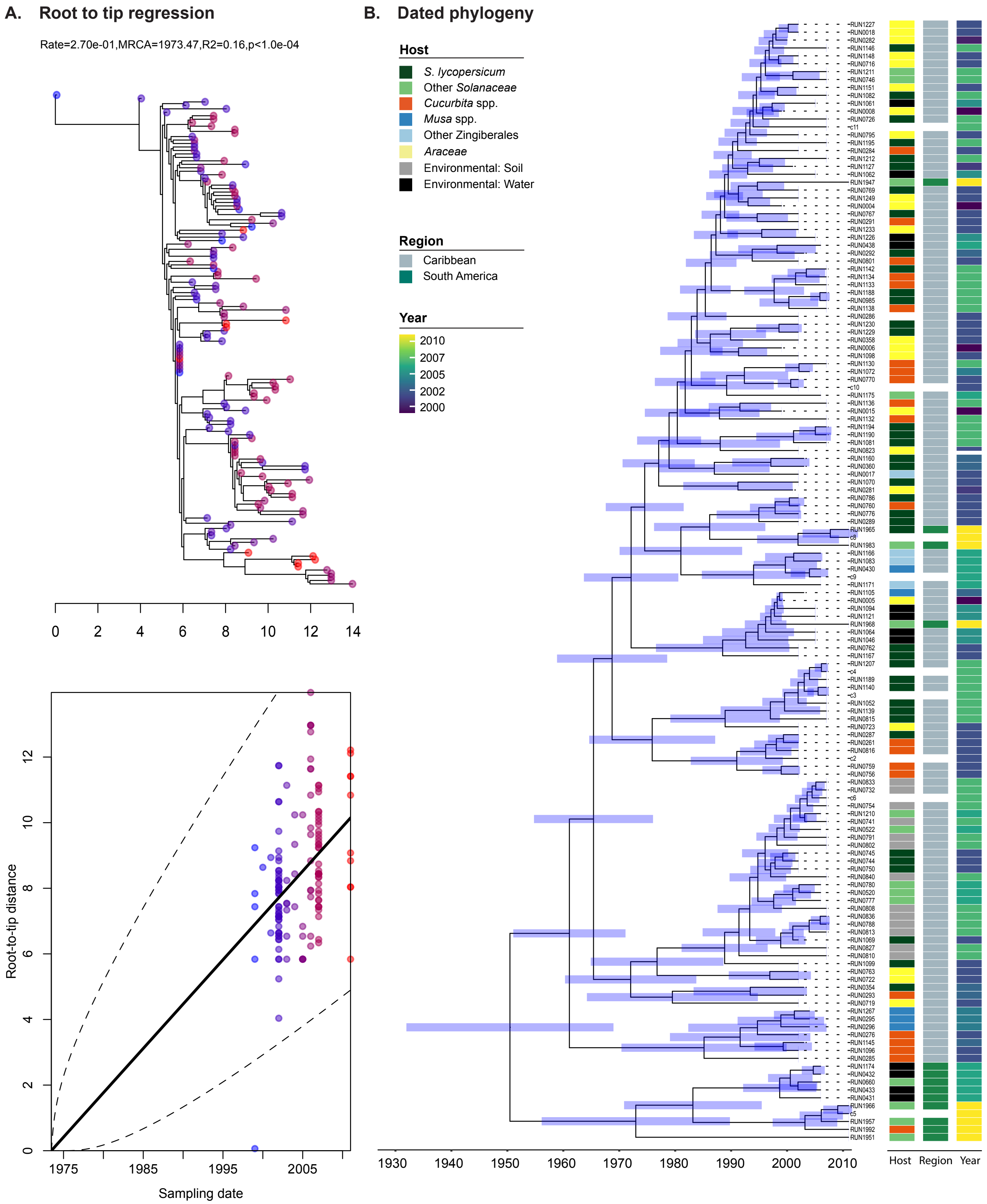

**Fig. S6. Divergence dating of Rso IIB-4NPB.** (A) BactDating root-to-tip regression plots for the Rso IIB-4NPB strains. (B). Maximum likelihood tree of all Rso IIB-4NPB with the temporal dating confidence intervals calculated by BactDating shown as blue horizontal lines. The clades collapsed due to zero-branch length are labelled Coll-Clade1 to Coll-Clade7. Coll-Clade1 contains strains RUN0727, RUN1193, RUN1192, RUN1104, RUN1137, RUN1075, RUN1151. Coll-Clade2 contains strains RUN0005, RUN0739, RUN0762, RUN1121, RUN1105, RUN1094, RUN1046, RUN1064. Coll-Clade3 contains strains RUN1953, RUN1980, RUN1973. Coll-Clade4 contains strains RUN0747, RUN0789, RUN0806, RUN0841, RUN0790, RUN0809, RUN0787, RUN0799, RUN0768. Coll-Clade5 contains strains RUN0788, RUN1069, RUN0836, RUN0813, RUN0827, RUN0840. Coll-Clade6 contains strains RUN1956, RUN1970, RUN1982, RUN1967. Coll-Clade7 contains strains RUN2004, RUN2011.

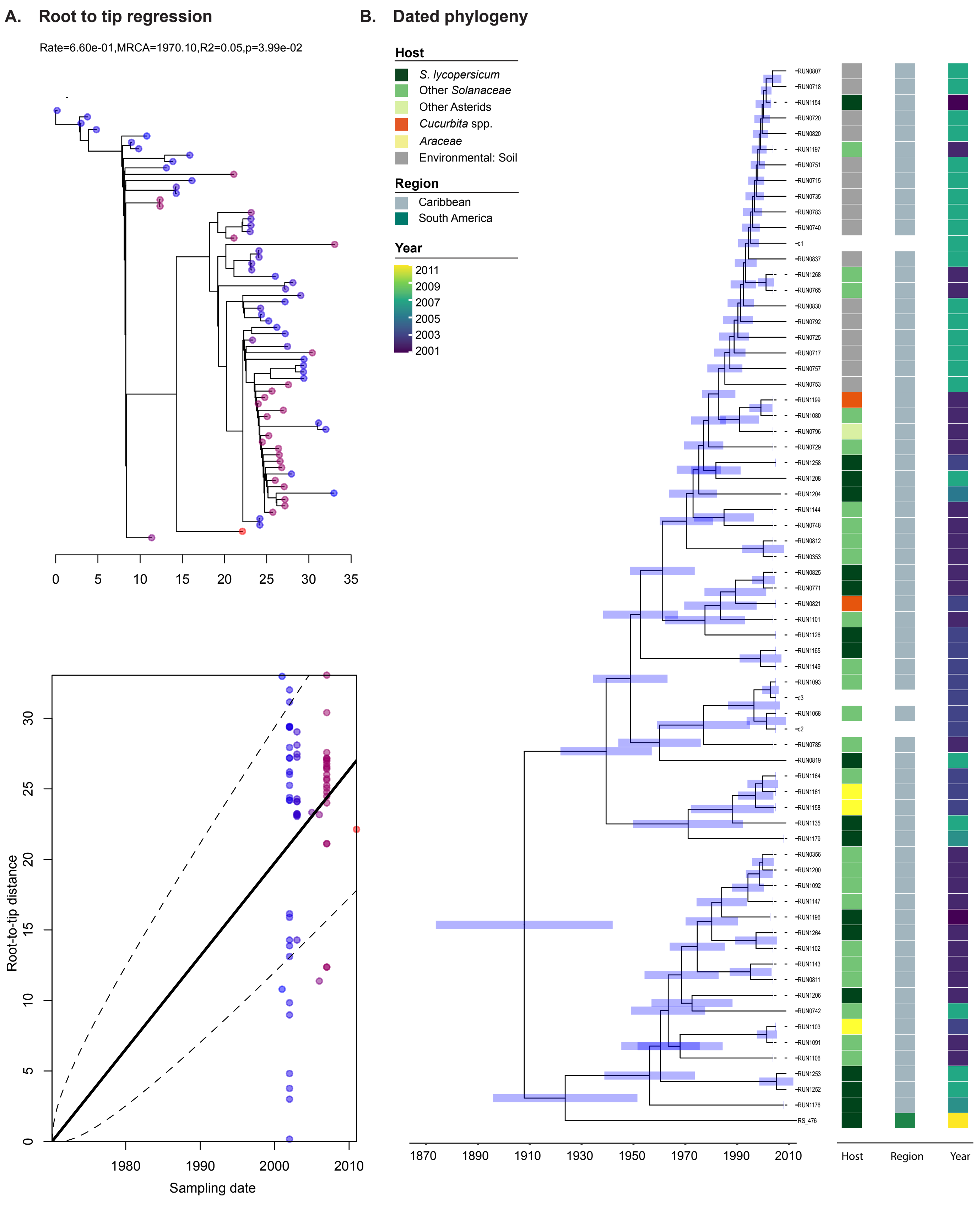

**Fig. S7. Divergence dating of Rps I-18.** (A) BactDating root-to-tip regression plots for the Rps I-18 strains. (B). Maximum likelihood tree of all Rps I-18 with the temporal dating confidence intervals calculated by BactDating shown as blue horizontal lines.

A Core genome tree

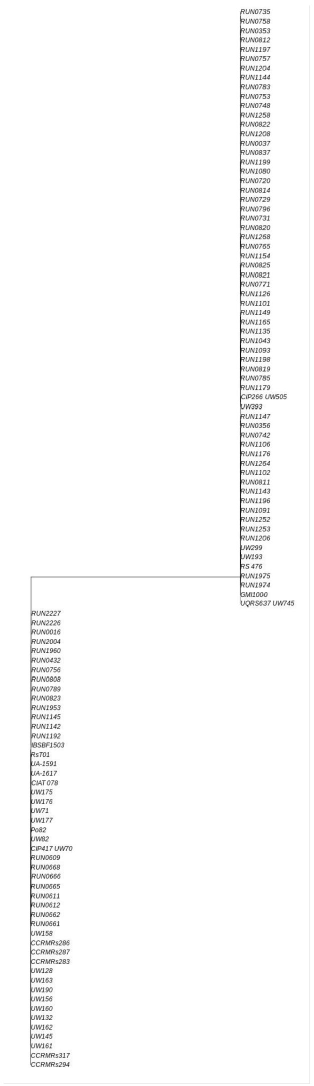

B Circularized core genome tree

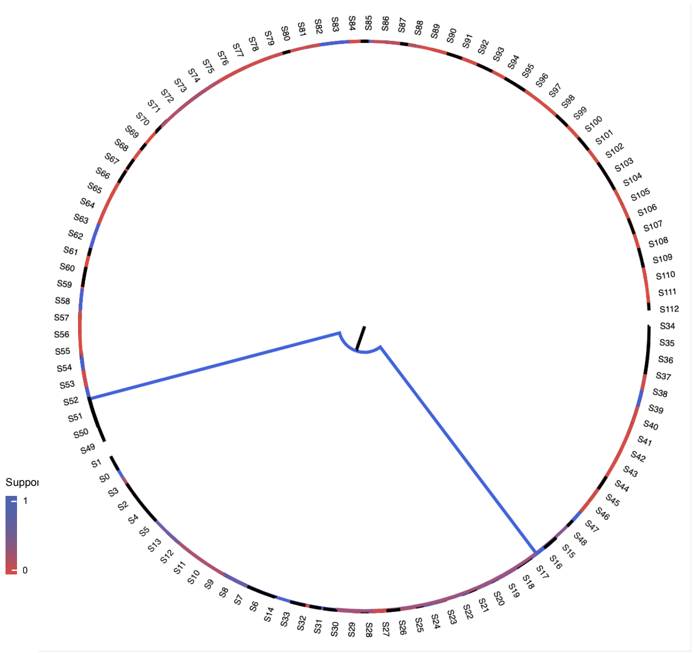

C SNPs per 1 KB block along the core alignment

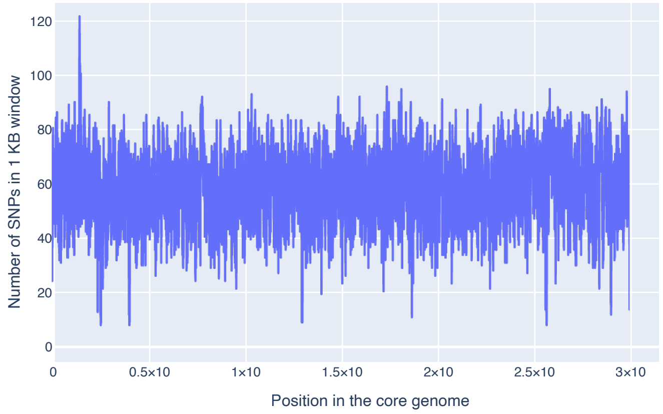

D Histogram of SNPs per 1 KB block along the core alignment

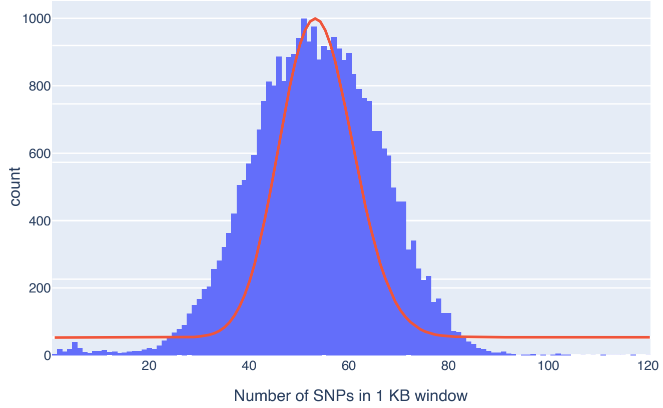

Diversity in core genome alignment

| | Invariant | Bi-allelic | Tri-allelic | Tetra-allelic |
|---|---|---|---|---|
| Numbers | 4,986,078 | 2,873 | 19 | 2 |
| Fractions | 0.999420 | 0.000580 | 0.000000 | 0.000000 |

**Fig. S8. RECOPHY analysis for reduced set of Rso IIB-4 and Rps I-18 genomes.**(A) Core genome tree generated by PhyML. (B) The same tree in circular form with the branches coloured according to the branch support values. (C) Plot representing SNPs counts per 1 kb of core alignment. (D) Histogram of SNPs counts per 1 kb.

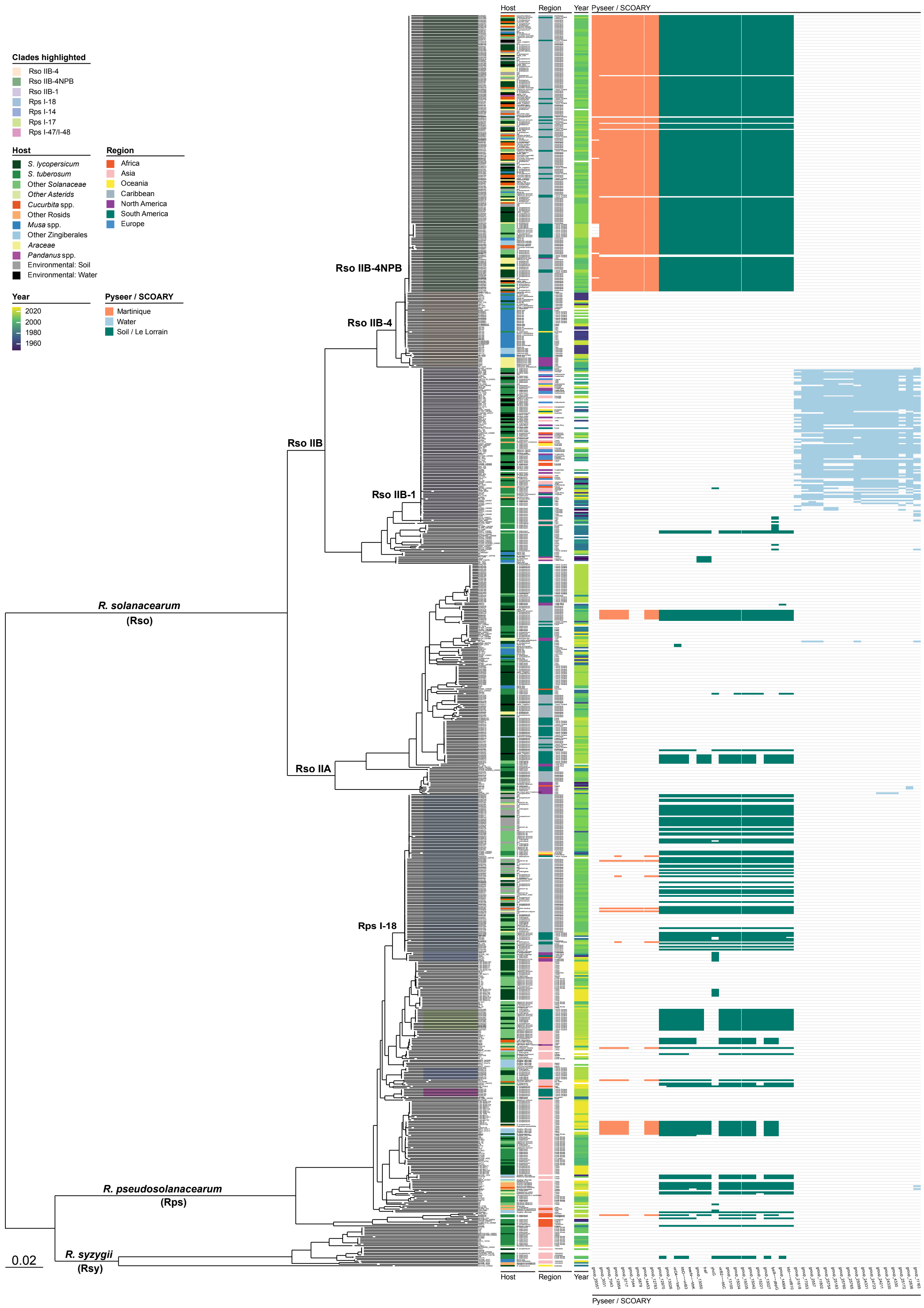

**Fig. S9. Complete panGWAS results displayed against RSSC core gene tree.** Maximum likelihood tree of all RSSC genomes with the metadata as described in Figure S1. Clades representing Rso and Rps lineages of interest are highlighted in the tree. Presence and absence patterns of all genes reported by both Scoary and Pyseer panGWAS analysis is shown on the right. The genes found in association with sampling in Martinique and Soil/Le Lorrain traits are shown in orange and green, respectively. The genes found in association with water samples are shown in blue.

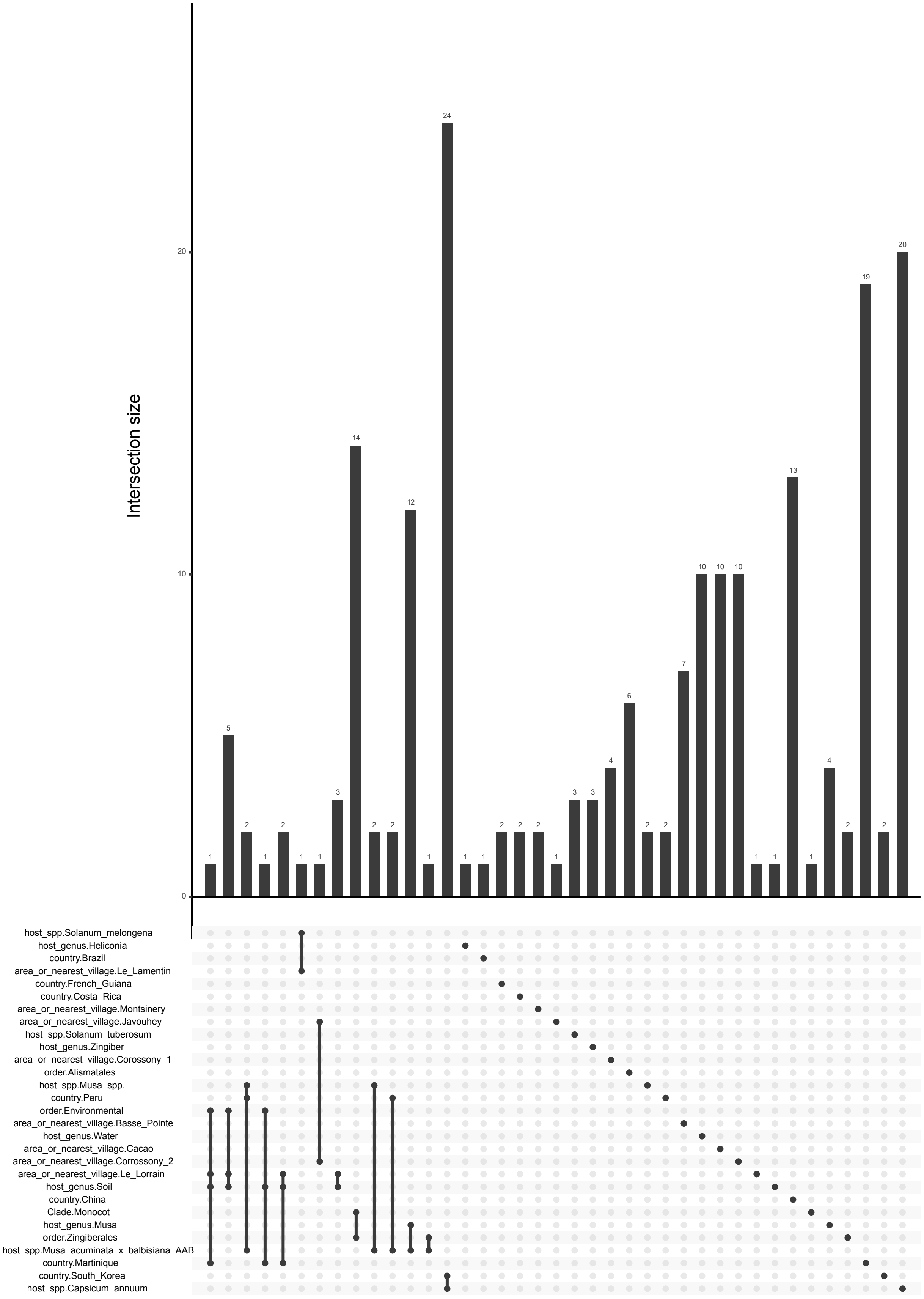

**Fig. S10. Pyseer UpSet plot.** Upset plot representing overlap between gene groups found in significant association with any of the traits tested by pangenome-based PySEER analysis. Rows show the region or host of isolation traits. Links in the plot indicate identical genes associated with 2+ traits. Bars represent counts of such genes.

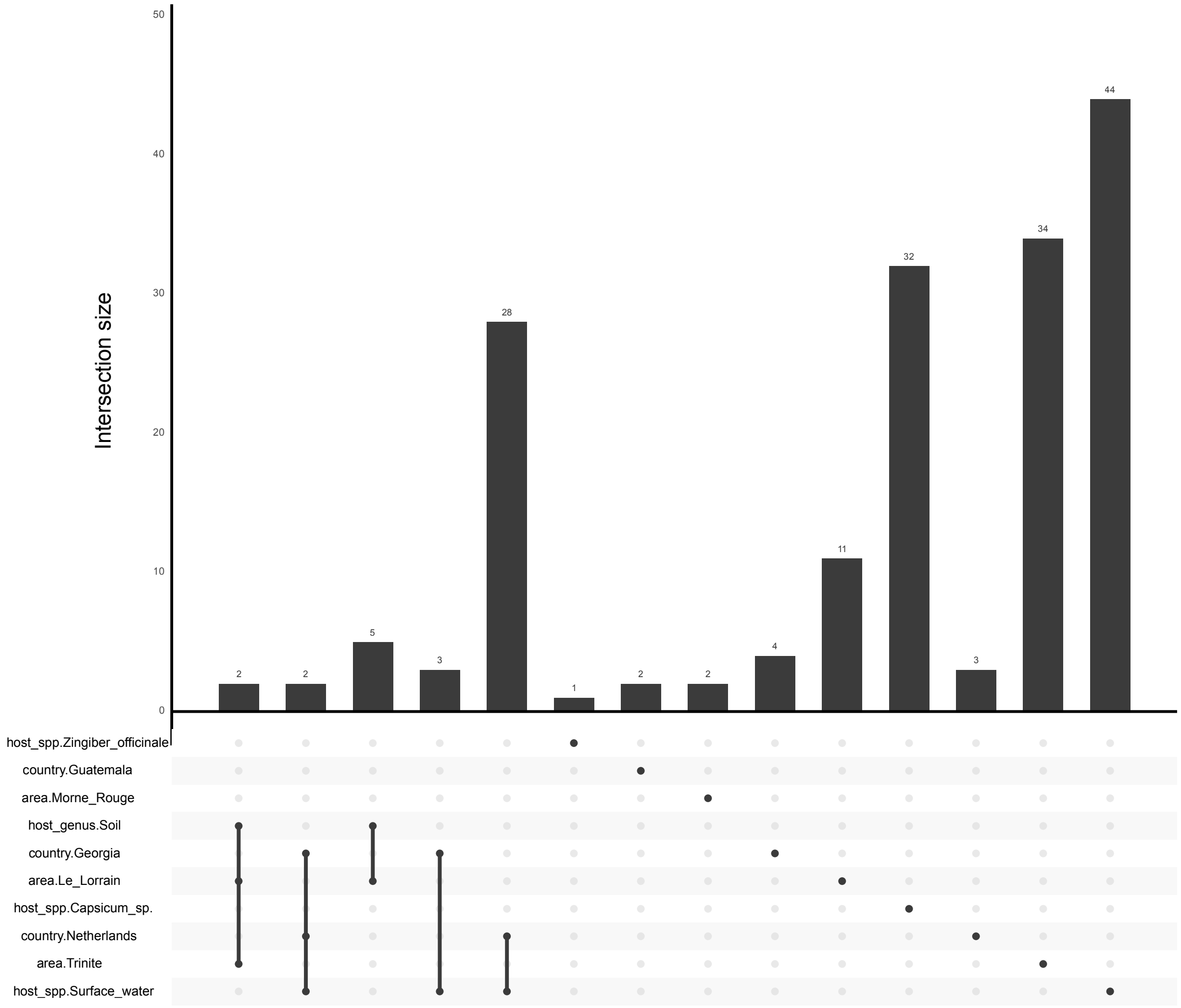

**Fig. S11. SCOARY UpSet plot.** Upset plot representing overlap between gene groups found in significant association with any of the traits tested by pangenome-based SCOARY analysis. Rows show the region or host of isolation traits. Links in the plot indicate identical genes associated with 2+ traits. Bars represent counts of such genes.

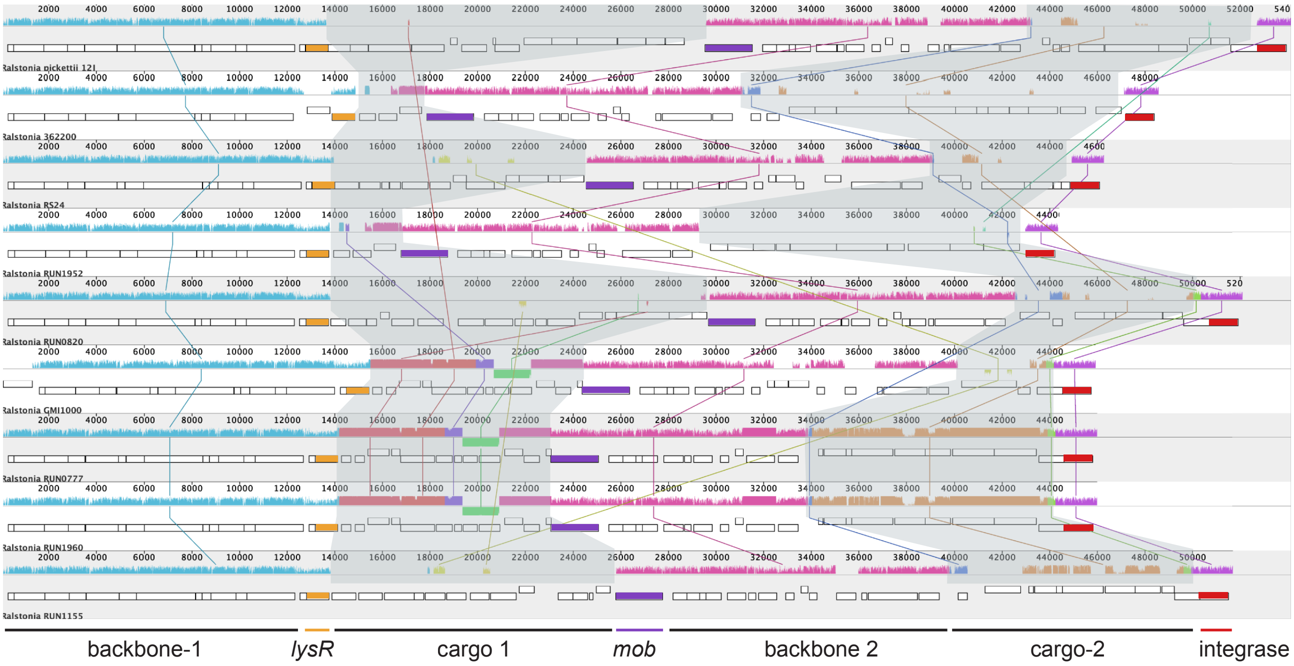

**Fig. S12. ICE synteny across Rso and Rps genomes.** ProgressiveMauve alignment of ICEs identified in *Ralstonia pickettii*, two Asian Rps strains (RS24 and 362200), three Rso and three Rps strains sampled in Martinique and French Guiana. The first backbone region (backbone-1) includes the T4SS and has a *lysR*-family transcriptional regulator at the 3' end, followed by a variable region called cargo-1. The second conserved region (backbone-2) spans the central 8kbp of the ICE and includes the *mob* relaxase, *traF*, *parA*, *repA*, *qseC* and *rdsF*, as well as genes encoding a helix-turn-helix domain-containing proten, a hypothetical and various DUF domain-containing proteins (DUF3800, DUF945, DUF2285 (*fseA*), DUF2840, DUF2958). A second variable region (cargo-2) is flanked by an integrase at the 3' end of the ICE. The blue and pink shaded similarity plots separately indicate the presence of backbone-1 and backbone-2 regions conserved across all ICEs, while the grey regions highlight the location of cargo-1 and cargo-2 regions. The lysR, mob and integrase genes are indicated separately as they delineate backbone and cargo region boundaries.

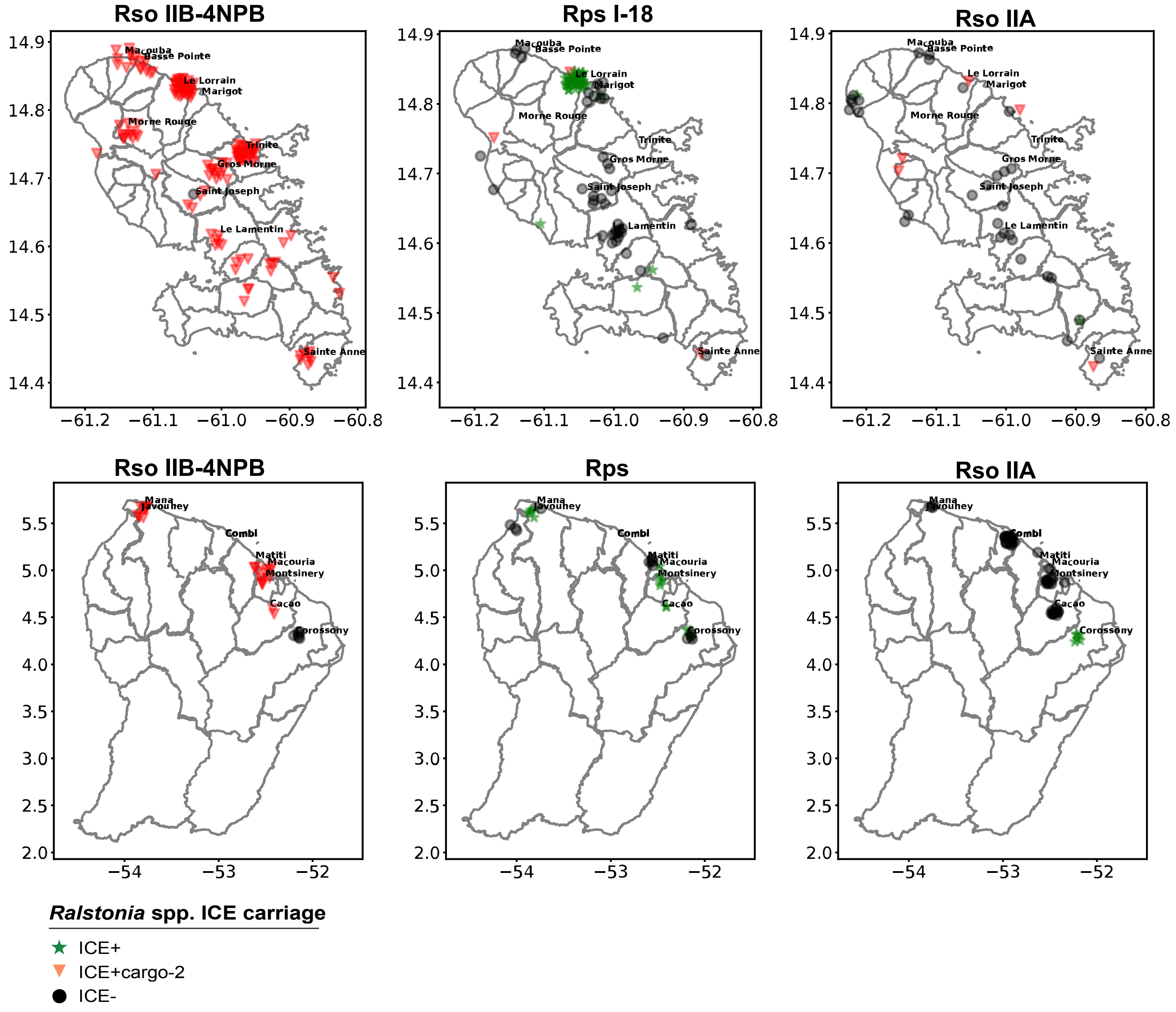

**Fig. S13. Geographic distribution of *Ralstonia* spp. ICEs sampled in Martinique.** Sampling sites of all strains collected in Martinique and French Guiana and assembled in this study. The exact longitude and latitude of the sampling locations were jittered for the visualisation purpose. The first and second row of the six panels show Martinique and French Guiana maps respectively with the locations where each Rso IIB-4NPB, Rps and Rso IIA strain was sampled. All Rps strains found on Martinique are Rps I-18, while Rps found in French Guiana belong to other lineages. Strains represented as red diamonds contain ICEs with cargo-2 regions. Strains represented as green stars contain ICEs without cargo-2. Black circles represent the strains without both insertions. Only the geographical locations where more than five strains were sampled are labelled with their names.

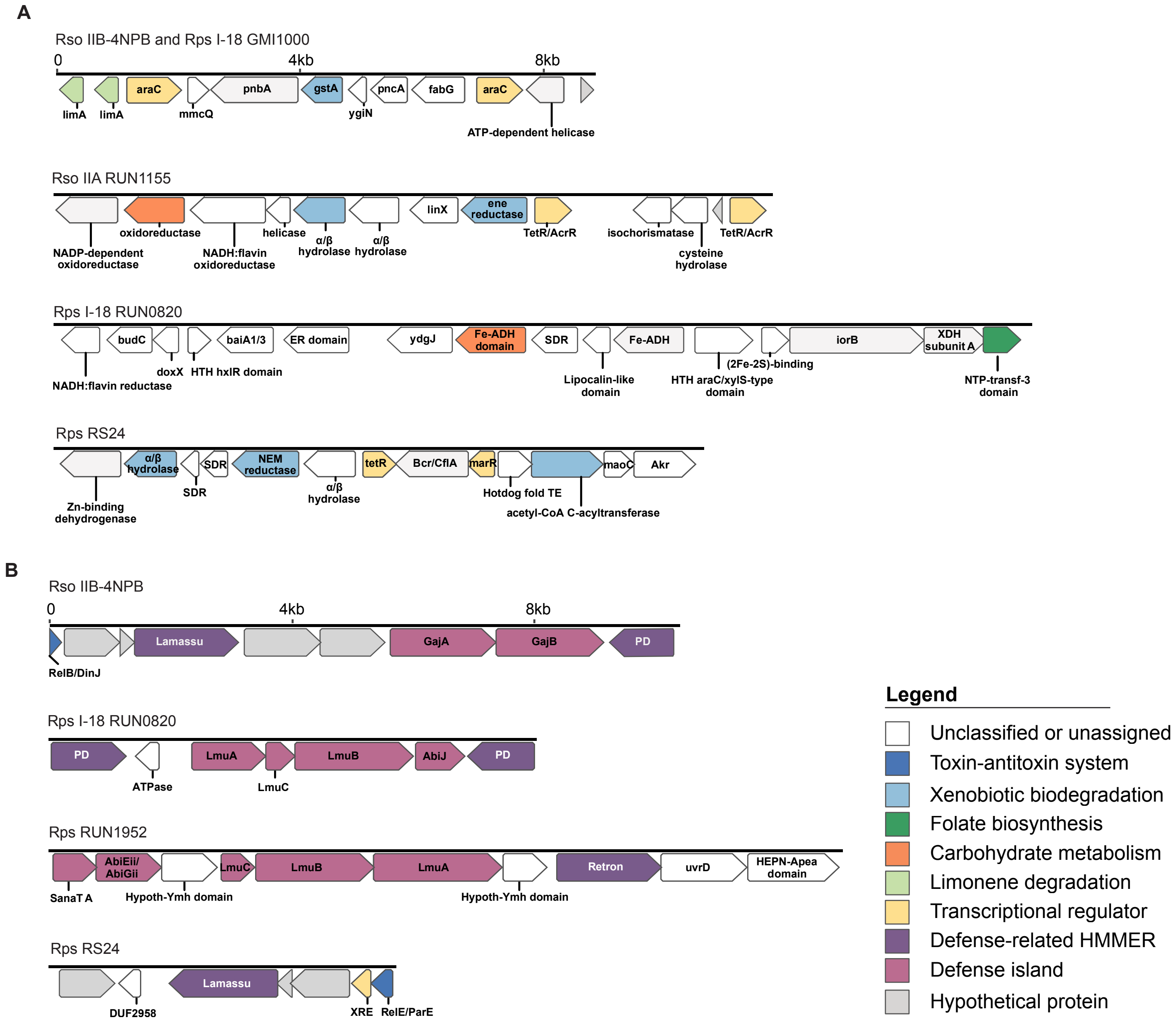

**Fig. S14. ICE cargo regions across multiple Rso and Rps strains.** (A) Functional annotations within cargo-1 regions of the ICEs found in all Rso IIB-4NPB strains, Rps I-18 GMI1000, Rso IIA RUN1155, Rps I-18 RUN0820 and Rps Rs24. Annotations of cargo-1 regions from Rso IIB-4NPB and Rps I-18 GMI1000 are identical and shown together. The CDS are coloured according to their KEGG annotation. Proteins assigned to a KO-term not classified as a part of a metabolic pathway are displayed in faint blue. Proteins identified as a part of xenobiotics biodegradation, folate biosynthesis, carbohydrate metabolism or limonene degradation pathways are shown in blue, dark blue, orange and red respectively. Transcriptional regulators and hypothetical proteins are shown in yellow and black respectively. (B) Functional annotations within cargo-2 regions found in all Rso IIB-4NPB strains, Rps I-18 RUN0820, Rps RUN1952 and Rps Rs24. CDS are coloured as in (a). Additionally, members of three toxin-antitoxin systems RelE/ParE, AbiEii/AbiGii and RelB/DinJ are shown in pink.

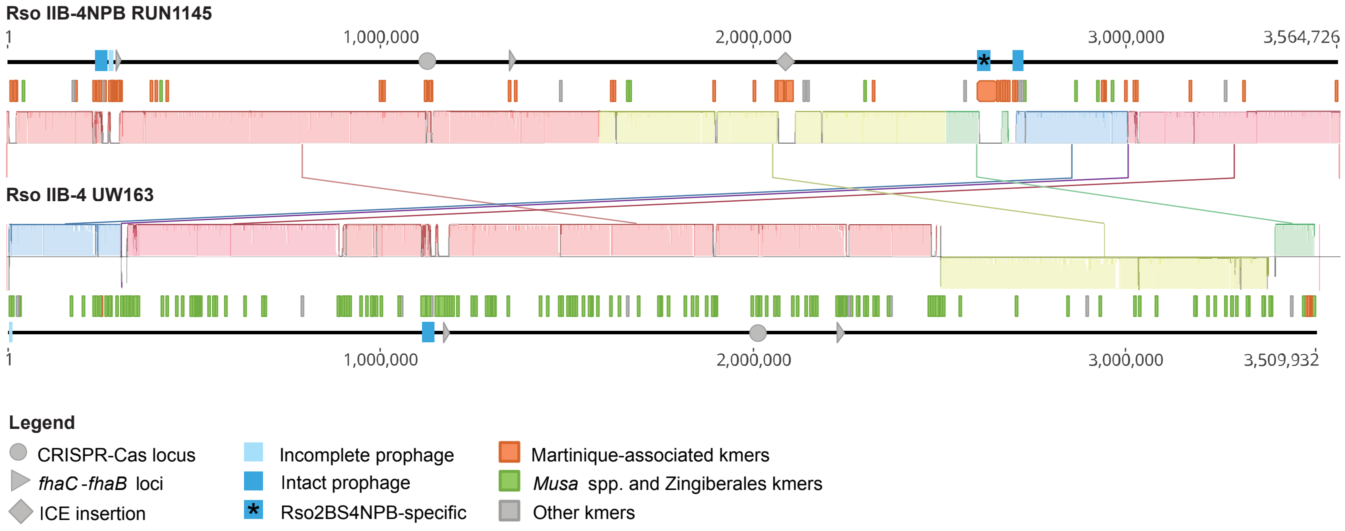

**Fig. S15. Mobile elements and host-associated kmers in Rso IIB-4 and Rso IIB-4NPB.** ProgressiveMauve alignment of the chromosomes of Rso IIB-4NPB RUN1145 and Rso IIB-4 UW163 strains shown in grayscale together with mapping positions of kmers reported by GWAS and annotated chromosome length scales. The alignment starts from the Rso IIB-4NPB RUN1145 origin of replication, while the Rso IIB-4 UW163 origin of replication is located at approximately 882,000bp. The chromosome length scales are annotated with positions of prophages (blue rectangles), CRISPR-Cas loci (grey circles), *fhaC-fhaB* loci (grey arrows) and the ICE insertion in Rso IIB-4 RUN1145 (grey diamond). The prophage uniquely found in Rso IIB-4NPB genomes is marked with an asterisk. The kmers that overlap or are located within 1kb of each other were merged. The kmer mapping positions (merged or not) are represented by coloured bars between chromosome scales and ProgressiveMauve locally colinear blocks. The kmers associated with isolation from Martinique are shown in orange, those associated with *Musa* spp. or Zingiberales isolation are shown in green. The kmers associated with any other trait are shown in grey.

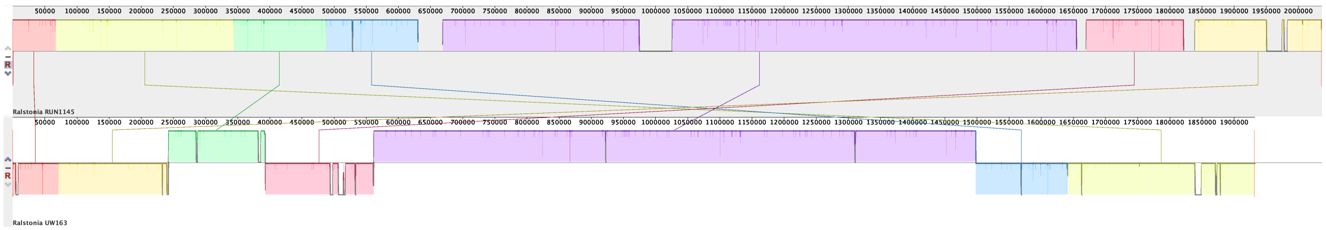

**Fig. S16. Alignment of Rso IIB-4NPB and Rso IIB-4 megaplasmids.** ProgressiveMauve alignment of the megaplasmids of Rso2BS4NPB RUN1145 and Rso IIB-4 UW163 strains.

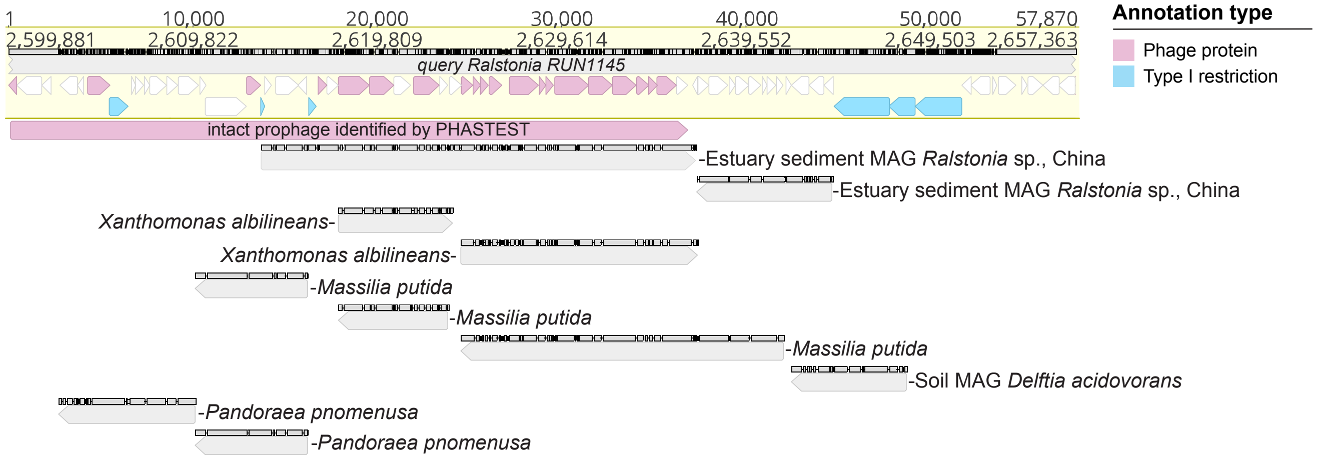

**Fig. S17. Rso IIB-4NPB-specific prophage BLAST results.** Prophage insertion at the 2,6-2,66Mb position within the Rso IIB-4NPB RUN1145 genome containing the phage unique to Rso IIB-4NPB. Pink arrows represent genes annotated as phage related: tail, head, phage terminase, and baseplate. Genes annotated as parts of the Type I restriction-modification system are shown in blue. The large pink arrow designates the position of an intact phage identified by PHASTER. Grey arrows below show hits reported by a BLAST search of the insertion region against the NCBI-nr database.

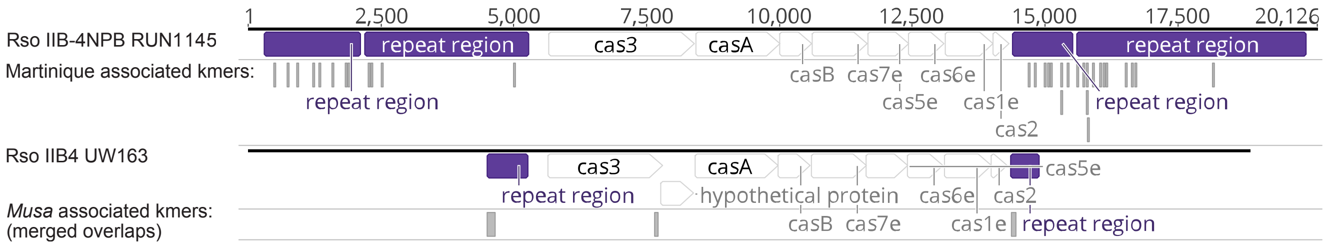

**Fig. S18. CRISPR-Cas loci in Rso IIB-4 and Rso IIB-4NPB.** GWAS kmers mapped to CRISPR-Cas loci located on the chromosome at the positions 1,119,872-1,139,675 and 2,010,416-2,020,801 in Rso IIB-4NPB RUN1145 and Rso IIB-4 sUW163 genomes, respectively. CRISPR-repeat regions are highlighted in purple. All kmers mapping to RUN1145 are associated with isolation from Martinique. All kmers aligned to UW163 are associated with isolation from *Musa* spp. Merged kmers are shown for those mapping to UW163, since many of these overlap. RUN1145 kmers are displayed without modifications.

### List of Supplementary Tables and Scripts

#### Tables

*Provided separately as an Excel file called Evseeva_et_al_2025_SuppTables.xlsx*

Table S1. RSSC genomes and associated metadata
Table S2. Panaroo pangenome gene presence and absence across RSSC
Table S3. Rso IIB-4-specific genes
Table S4. Rso IIB-4NPB-specific genes
Table S5. Rps I-18-specific genes
Table S6. T3E family distribution across the RSSC
Table S7. Phages identified in 17 complete RSSC genomes
Table S8. Rso IIB-4NPB CRISPR spacer BLAST hits
Table S9. Pyseer kmer-based trait associations
Table S10. Pyseer kmer mapping positions
Table S11. Pyseer panGWAS results
Table S12. SCOARY panGWAS results
Table S13. MOBP relaxase positions identified by MOBscan
Table S14. Assembly data for all genomes acquired from NCBI

#### Scripts

*Scripts are provided separately as a zip file called Evseeva_et_al_2025_python_scripts.zip*

kmers_map2ref.py
presence-absence-tables.py
pyseer_upset.py
scoary_ovelaps_n_filter.py
t4ss_blast_ICE_pos_to_table.py

### Supplementary Results

#### Identification of ICEs in the *Ralstonia solanacearum* species complex

All eight $ICE^{RpsGMI1000}$ identified in Rps genomes have integrated into one of three attachment (att) sites on the primary chromosome, numbered incrementally in order of distance from the origin of replication in Rps I-18 GMI1000 (i.e. att-1, att-2 and att-3). Two Rps genomes have $ICE^{RpsGMI1000}$ in att-1, four Rps genomes have $ICE^{RpsGMI1000}$ in att-2, and three genomes have $ICE^{RpsGMI1000}$ in att-3. ICE attachment sites frequently overlap with tRNA genes, but this is not the case here, and comparison of the flanking regions shows structural rearrangements have occurred[1]. The att-1 site appears least preferred for ICE integration, and is located between genes encoding a hypothetical protein and a purine metabolism enzyme phosphoribosylformylglycinamidine cyclo-ligase. The att-2 is located between XRE family transcriptional regulator and inositol monophosphatase genes, while att-3 is located between genes encoding a universal stress protein and isocitrate lyase. All five ICEs identified in complete Rso assemblies are $ICE^{RpsGMI1000}$ integrated into att-2 sites in Martinique isolates, although two Rso IIB-4NPB draft assemblies (RUN1960 and RUN1959) contain an ICE integrated into the att-3 site. The att-3 is the preferred integration site in Rps genomes for most complete or partial ICEs, $ICE^{RpsGMI1000}$ or otherwise.

Rps ICEs retain core backbone genes (*trb* conjugative system, *traG* coupling protein, *mob* relaxase and integrase) but vary considerably in cargo, i.e. gene clusters that do not encode functions critical to the ICE lifecycle, but are highly variable between ICEs and may confer novel fitness-enhancing functions to the bacterial host. Conserved and cargo regions in the broader family of ICEs were delineated using ProgressiveMauve alignment of complete ICEs from five Rps genomes representing five clades in the phylogenetic tree, three Rso (Rso IIA and two Rso IIB-4NPB with ICEs in either att-2 or att-3) and one outgroup *R. picketii* genome (Fig. 4). The cargo-1 in Rps I-18 RUN0820 and Rps RUN1155 ICEs encodes genes associated with carbohydrate metabolism and folate biosynthesis[2], and Rps RUN0820 ICE cargo-1 encodes various reductases, oxidases and dehydrogenases involved in the metabolism of fucose, a sugar found on plant cell surface (gene ID PEIJFM_11570)[3].

#### Virulence gene repertoires in the *Ralstonia solanacearum* species complex

Patterns of T3E family presence and absence correlate with genetic distance, i.e. strains within the same species or the same clade share similar T3E repertoires. The average pairwise similarity within T3E families is higher within Rso (92%) and Rps (94%) than it is across the entire species complex (86%). *R. syzygii* (Rsy) exhibits the highest diversity among T3E families (88%), despite being less well sampled than both Rso and Rps. The core effector arsenal of the species complex is comprised of 31 T3E families present in more than 99% of all RSSC genomes: *ripA3, ripA4, ripA5, ripB, ripC1, ripD, ripE1, ripG4, ripG7, ripH2, ripM, ripR, ripS2, ripS5, ripU, ripW, ripY, ripZ, ripAB, ripAC, ripAD, ripAE, ripAI, ripAJ, ripAM, ripAN, ripAO, ripAP, ripAR, ripAY, ripTPS*. Some T3Es were found in only a single RSSC genome: RipO2 is present solely within *Rsy* strain NCPPB_3219 from clove (*Syzygium aromaticum*), RipBN is limited to Rps3S29 strain CMR15 from tomato (*S. lycopersicum*) and the hypothetical effector Hyp17 is present solely in Rps strain Tb04 from tobacco (*Nicotiana tabacum*) (Fig. S1). Five T3E families were not identified in any of the RSSC genomes despite being listed in the RalstoT3 database (*ripG8*, *ripAF2*, *ripBE*, *ripBP* and *ripBQ*). This may be due to filtering and assignment to a T3E family based on similarity of the best hit, or the exclusion of the relevant genomes from our dataset. A single Australian isolate grouping with the Rso IIB-4 strains carrying *ripP3* was also isolated from *S. tuberosum*.

Rps I-18 displays a uniform effector repertoire, with one exception: *ripS4* is variable across Rps I-18. The gain and loss of *ripS4* does not appear to be linked with host, substrate or location of isolation however. The same effector is often missing in other Rps clades circulating in French Guiana, but present in the majority of Rps strains from Asia. Rps I-18 also lacks *ripAL*, an effector otherwise present in almost every other RSSC genome. Conversely, Rps I-18 acquired *ripBO*, an effector almost entirely absent from every sequenced Rps and Rso (Fig. 3). The effector *ripAX2* has been demonstrated to trigger specific resistance in eggplant[4].

While there is no evidence variation in T3E carriage is linked with the emergence and host range expansion of Rso IIB-4NPB, T3E exchange may play a role in Rso IIB-4 pathogenicity on banana. The Rso IIB-4 isolates from cultivated endemic (*Heliconia* spp.) and introduced (banana, *Epipremnum* spp.) monocot hosts that are endemic and introduced (banana and) to the Americas exhibit more variation in effector repertoire than Rso IIB-4NPB (Fig. 3, Fig. S2). Two effectors are present in Rso IIB-4 and absent in Rso IIB-4NPB: *ripAX2* and *ripP3*. The *ripAX2* gene is present in many other Rso IIB and Rso IIA clades, indicating loss in Rso IIB-4NPB is likely, while ripP3 is sparsely distributed in Rso IIB. Four effector families exhibit evidence of likely loss in monocot-infecting Rso IIB-4: ripAQ, ripAW, ripBD and the predicted T3E_Hyp3 effector. The loss of ripAQ is most striking as the family is otherwise present in all other RSSC strains. The ripAW gene is missing from most Rso IIB-4 banana isolates from Peru, most pandemic Rso IIB-1 potato isolates and some R. syzygii, but is present in all other RSSC strains. Peruvian Rso IIB-4 strains also lack ripBD, an effector otherwise common in Rso IIA. T3E_Hyp3 are absent from Rso IIB-4 and patchily distributed in other Rso IIB lineages. Since these effector families are present among related clades and/or across the species complex, the most parsimonious explanation for the pattern observed is they were lost in the ancestor of banana-infecting Rso IIB-4 strains rather than being acquired by ancestral Rso IIB-4NPB. Their link with the pathogenicity of Rso IIB-4 on monocots *Musa* spp., *Heliconia* spp. and *Epipremnum* spp. is not known.

**Genomic variants associated with host of isolation**

Guided by the possibility the depth of sampling may allow us to uncover variants associated with isolation from a particular host, we applied three methods for genome-wide association testing. We took advantage of the extensive metadata available to run genome-wide tests for associations between genetic variation (either genes or kmers), host and location of isolation.

Among 80,586 kmers reported by Pyseer, many kmers (22,665) were linked with specific hosts, most frequently Musa spp. (8,229 kmers), Heliconia spp. (4,560 kmers), Capsicum annuum (2,554 kmers) and members of the Alismatales (Anthurium spp., Epipremnum spp., and Pothos spp., 2,369 cumulative kmers) (Table S4). Mapping kmers against Rso IIB-4NPB RUN1145 showed that its genome contains 36 Musa-associated kmers in 16 loci including a region containing FAD-binding oxidoreductase (OFBKGH_00185), alpha/beta hydrolase (OFBKGH_00190), membrane protein (OFBKGH_00195) and response regulator transcription factor (OFBKGH_00200 in RUN1145, Table

S10). Remaining Musa-associated kmers map to insertion sequences, two prophages, multiple intergenic regions, hypothetical protein-encoding genes, glycine hydroxymethyltransferase (OFBKGH_12725 in RUN1145, Table S10). 52.6% (2,316/4,401) of kmers mapping to the Rso IIB-4 UW163 reference genome are associated with isolation from banana. In comparison, only two banana-associated kmers map to the RpsS18 GMI1000 genome.

The *Musa* associated variants identified in the strain the Rso IIB-4 UW163 are found in various genes including several Type III effector proteins: UW163_05335 (*ripP3*), UW163_14335 (*ripV2*), UW163_20305 (*ripS3*), UW163_20590 (*ripA5*), UW163_20945 (*ripC1*) and UW163_22160 (*ripAT*). Recombination breaks up linkage disequilibrium, and the repeated independent evolution of pathogenicity on banana across the species complex increases the likelihood of identifying variants linked with convergent trait evolution. These finding also align with the suggestion pathogenicity on this host is a derived trait in Rso IIB-4[5–7].

The emergence of a pandemic lineage infecting crop plants is frequently linked with the introduction and expanded cultivation of susceptible host varieties[8–11]. The center of origin of *R. solanacearum* is in the Americas, where banana was introduced as a crop five centuries ago and became intensively cultivated only in the last two centuries[12]. Infectious disease outbreaks occurred soon after, most notoriously those caused by *Fusarium oxysporum* TR1 and *F. odoratissimum* TR4, which previously eliminated the Gros Michel variety and now threaten the Cavendish variety[13,14]. The banana-associated genes and kmers identified here represent interesting candidates for study of the genetic basis of specialisation on this specific host. Although further work is required to determine whether the candidates identified in this work are responsible for banana pathogenicity in Rso and the increased fitness of Rso IIB-4NPB in soil and host environments, this work shows bacterial metabolism is likely to play an important role in host adaptation, and reveals how ecological interactions with other members of the microbial community shape pathogen evolution and contribute to the emergence of infectious disease.

Gene-based panGWAS methods (SCOARY and Pyseer) were also applied to search for genes whose presence or absence is associated with a particular host of isolation. Pyseer reported a total of 938 genes significantly associated with any region or host of isolation traits (Table S11). Among these, multiple genes associated with isolation from *Musa* spp. (135), *Solanum melongena* (80), *Capsicum annuum* (56) and *Heliconia* spp. (21) were identified. There was no overlap between the Pyseer and SCOARY predictions for host of isolation, though there were for association with specific locations, at varying spatial scales (Table S12).

**Spatial distribution of genetic variation**

There are many genes found solely in association with specific location, at varying spatial scales (Table S12). The overlapping set of genes identified by both SCOARY and Pyseer panGWAS includes 27 genes associated with isolation from specific districts (communes) in Martinique (Fig. S10, Table S11, Table S12). Many kmers (57,921) were identified as being associated with a specific district (commune) or country (Table S9). Very few had overlapping geographic associations (for example between country and district), but 32% of these had overlapping associations between location and host of isolation. For example, some kmers show evidence of association with isolation from both banana and Peru. Mapping kmers against reference genomes shows there is ample genetic variation associated with Martinique among both Rso IIB-4NPB and Rps I-18. This is not unique to Martinique: there are also China and Peru-specific kmer associations (Table S10). 93.5% of the kmers mapping to Rso IIB-4NPB RUN1145 are associated with isolation in Martinique.

As mentioned, the majority of location-specific genes identified by SCOARY and both Pyseer methods are located on the ICE$^{RpsGMI1000}$. Outside mobile elements, the most common genes with location-specific associations are fhaC and fhaB (Fig. 5). These encode type 5b two partner secretion systems comprised of a transporter (FhaC) and extracellular hemagglutinin repeat-containing protein (FhaB). Interestingly, kmers associated with isolation from two distinct regions can map to different regions of the same locus. For example, three out of seven fhaC-fhaB loci in Rps I-18 GMI1000 have China-associated kmers mapping to the 5' end of fhaB and Martinique-associated kmers mapping to the 3' end of the same gene (GMI1000_04540, GMI1000_09170 and GMI1000_20245). The 3' end of another fhaB gene is composed of alternating regions containing either China or Martinique-associated kmers mapping (GMI1000_22860). Fine-scale spatial associations with fhaC-fhaB are also present, for example between an fhaB and the region of Le Lorrain in Martinique. Overlap with predictions for host association occurs as well: kmers associated with isolation from both *Musa* spp. and Peru map to four of six fhaC-fhaB loci from Rso IIB-4 UW163 (UW163_05665, UW163_10515, UW163_21610 and UW163_24515).

**CRISPR-Cas loci in Rso IIB-4 and Rso IIB-4NPB**

The CRISPR-Cas operon (*cas3*, *casA/cas8e*, *casB/cas11/cse2*, *cas7e*, *cas5e*, *cas6e*, *cas1e*, *cas2*) is conserved between Rso IIB-4-UW163 and Rso IIB-4NPB RUN1145 (a single nucleotide frameshift mutation resulting in a premature stop in Rso IIB-4 UW163 *cas3* is likely a sequencing error as it is located in a region of 6 consecutive cytosines). The kmer-based association tests also show that the largest cluster of Martinique-associated kmers that do not map to MGEs map to the leader-proximal position of CRISPR arrays in Rso IIB-4NPB. The 31bp kmers partially overlap with the near identical 29bp CRISPR 1 and CRISPR 2 repeat sequences, as well as up to 7bp of spacer sequence, with CRISPR repeats sometimes differing in the final 1-2bp.

The INPHARED phage database was queried using Rso IIB-4 and Rso IIB-4NPB spacers, yielding hits to nine different Ralstonia phages: RPZH3[15], RS603, RSMSuper[11], Eline, Raharianne[16], YO010_2, YO134_2, YO026_1 and YO158_1[17] (Table S8). Spacers shared between CRISPR arrays of both Rso IIB-4NPB RUN1145 and Rso IIB-4 UW163 map to YO134_2, YO026_1 and YO158_1 phage. The other spacers with hits in the INPHARED database are among the set unique to and recently acquired by Rso IIB-4NPB RUN1145. None of the Rso IIB-4NPB RUN1145 spacers aligned to the RefSeq Phage or env_nt databases. NCBI-nr BLASTn reported two hits for Rso IIB-4NPB unique spacers: Rps3S23 (LN899820.1) and *Burkholderia* sp. S-53 (CP090483 and CP090484) genomes.
